# Artificial hibernation uncovers distinct synaptic engram architecture for memory retention

**DOI:** 10.64898/2025.12.09.692927

**Authors:** YJ Lin, A Takahashi-Nakazato, K Tsutsumi, T Takahashi, D Mercier, H Ashitomi, MC Chiang, M Haberl, M Uytiepo, A Maximov, Y Makino, T Nemoto, R Enoki, A Hirano, K Soga, S Looprasertkul, N Ohno, Y Kubota, T Sakurai, KZ Tanaka

**Author notes:** These authors contributed equally (YJ Lin, A Takahashi-Nakazato, K Tsutsumi, T Takahashi).

## Abstract

The memory trace at the neuronal and synaptic levels remains controversial. Stable, larger spines are thought to support memory, but the high turnover of dendritic spines and the drifting of neuronal representations following memory formation suggest alternative possibilities. To elucidate a structural trace underlying memory retention, we utilize a mouse model of artificial hibernation. During hibernation, hippocampal neurons exhibited a substantial reduction in their activity and an extensive elimination of dendritic spines and synapses. Despite these changes, their memory and associated hippocampal neuronal representations are intact after arousal. We find that a subset of spines is maintained during hibernation. These spatially clustered engram-engram synapses are exclusively protected from elimination and characterized by synaptic contacts with multi-synaptic boutons. These findings suggest that synaptic engram architecture, rather than larger spines per se, is resilient to network remodeling and underlies long-term memory retention.

## Main Text

Retention of memory requires the storage of a memory trace in the brain (*1–6*). Various types of experience-dependent changes in the brain play indispensable roles in the successful acquisition of memory (*7–12*). Among these changes, the long-lasting enhancement of synaptic efficacy, which is strongly correlated with spine volume, is widely accepted as a structural memory trace (*13–18*). Despite the prevailing expectation that neuronal underpinnings of memory are stable across the retention interval, many studies have reported high instability in dendritic spines and the drift of neuronal representations over days, especially in the hippocampus (*19–24*). This discrepancy raises many unaddressed questions. Do experience-dependent changes created during memory encoding must persist to sustain long-lasting memory? Are there key features retained in the face of overall instability that underlie memory retention? Recent studies of memory engram are also controversial in this regard. While enhanced connections are observed between engram cells, studies demonstrated that engram activation is sufficient to produce memory recall, even when strengthened synaptic connections are compromised and when memory is no longer recalled from sensory cues (*25*, *26*). These studies suggest that a particular network architecture, rather than the retention of individual strengthened synapses, underlies memory storage. Despite the fundamental significance of this problem, the complexity and dynamic nature of the neuronal network make it challenging to uncover the structural underpinning that achieves memory retention.

Hibernation may provide crucial insights into the structural mechanisms underlying memory retention. Past studies reported that hibernating mammals exhibit reduced neuronal activity and dendritic shrinkage in the hippocampus, suggesting that a metabolic energy-saving mechanism minimizes functions in the hibernating brain (*27–34*). Multiple species of hibernating mammals are reported to retain intact memory after arousal, despite the brain’s dormant state and reduced activity during winter (*35–37*) (but also see (*38–40*)). Structural changes in their brains during hibernation may offer a novel approach to minimize the complexity of the neuronal network without compromising memory, and to uncover the structural traces sufficient for memory retention. However, the use of natural hibernating mammals is hampered by difficulties in controlling experimental parameters and by limited methodologies for examining and manipulating the brain system. This study circumvented these limitations by harnessing artificial hibernation in mice.

## Results

### Extreme downscaling of the neuronal network during hibernation

We examined the mechanisms of memory in the hibernating brain using a mouse model of artificial hibernation (Q-neuron-induced hypometabolism and hypothermia, QIH; (*41*, *42*)). Chemogenetic activation of Q-neurons in AVPe (the anteroventral periventricular nucleus) produces a rapid and substantial reduction in body temperature that persists for approximately 48 hours (Fig. 1A). Single-unit tetrode recording from the dorsal hippocampal CA1 of the hibernating mice confirmed a rapid reduction of the firing rates after CNO injection (Fig. 1B, Fig. S1). In parallel, we observed a global suppression of LFP (local field potential) amplitude across broad frequency bands (Fig. 1C & D). Significant suppression of oscillations was observed in the theta, ripple, and low/high gamma frequency ranges (Fig. 1E). These observations indicate that a substantial reduction in neuronal activity accompanies metabolic suppression during QIH.

**Fig. 1.**
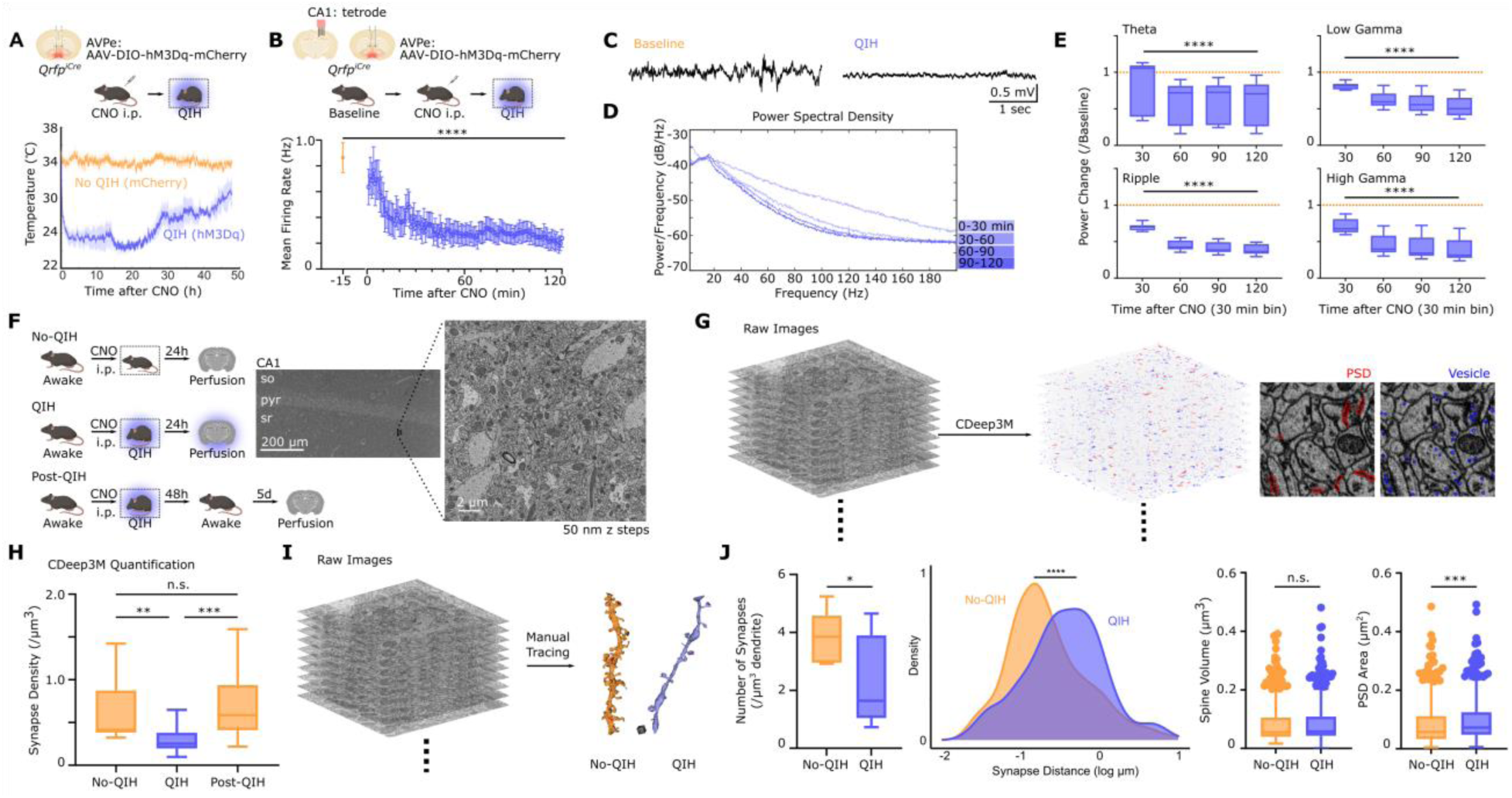
During a controlled hypothermia and hypometabolism (QIH) neuronal activity is reduced and dendritic spines are eliminated in the hippocampus. (A) Changes in the body temperature after CNO injection to the Qrfp-iCre mice with AAV-DIO-hM3Dq-mCherry expression in AVPe (N = 3 mice). Plots show the temperature values averaged over 2-minute bins and the standard errors of the means. (B) The firing rate of the the CA1 pyramidal cells is reduced after QIH induction (n = 155 units from 3 mice, Friedman test, p < 0.0001). Firing rates are averaged during the 15 minutes of the resting session (Baseline) or over 1-minute bins after CNO injection. Error bars indicate the standard error of the means. (C) Representative LFP trace from Baseline (left) or QIH (right) session. (D) Representative power spectral density plots from four 30-minute time bins after CNO injection. (E) Changes in the oscillation power after CNO injection (n = 12 recording channels in the hippocampus from 3 mice). Theta (4-12 Hz, p < 0.0001), low gamma (30-50 Hz, p < 0.0001), high gamma (65-85 Hz, p < 0.0001), and ripple (100-130 Hz, p < 0.0001) ranges were examined (Friedman test). The mean power in the baseline session normalizes power values. (F) Brain samples from three groups of mice are collected for volume EM (SBEM) imaging (No-QIH, QIH, Post-QIH; N = 3 mice per group). Right, representative EM images indicating an ROI at the stratum radiatum (sr). (G) A representative stack of raw EM images processed for CDeep3M detection of the post-synaptic densities (PSDs) and pre-synaptic vesicles (vesicles). PSDs or vesicles are highlighted in red or blue, respectively (right). (H) Synaptic density within an ROI (850 µm^3^) compared across the three groups (n = 16∼24 ROIs from 3 mice in each group, Kruskal-Wallis test followed by post-hoc multiple comparisons with Dunn’s correction, p = 0.0012 for No-QIH vs QIH, p = 0.0001 for QIH vs Post-QIH, p > 0.9999 for No-QIH vs Post-QIH). (I) A representative stack of raw EM images with neuronal membranes manually segmented from No-QIH and QIH animals (left) and representative dendrites 3D reconstructed from the manual segmentation (right). A gray cube indicates 1 μ m^3^ scale. (J) The number of synapses per unit volume of a dendrite was compared between No-QIH and QIH mice (left) (n = 7∼9 dendrites in each group, unpaired t-test, p = 0.0424). The distance of each synapse to the closest synapse on the same dendrite was compared between the two groups (middle) (n = 360∼362 synapses in each group, Kolmogorov-Smirnov test, p < 0.0001). The spine volume was compared between the two groups (n = 425-451 spines in each group, Kolmogorov-Smirnov test, p = 0.1313). The PSD area was compared between the two groups (n = 425-451 spines in each group, Kolmogorov-Smirnov test, p = 0.0001). Box plots show median, 1st and 3rd quantiles, minimum & maximum values within 1.5 times the interquantile range (IQR) from each quantile, and the other data points outside of the range.

We next examined structural changes of the hippocampal neurons during QIH. To this end, SBEM (Serial Block Face-Scanning Electron Microscopy) images of the hippocampal CA1 stratum radiatum in three groups of animals (No-QIH, QIH, and Post-QIH) were acquired to characterize their synaptic connections as well as ultrastructural features (Fig. 1F). The SBEM images were first processed with AI-assisted segmentation CDeep3M (*43*) to detect post-synaptic densities (PSDs) and pre-synaptic vesicles (Fig. 1G). Quantification of the synaptic contacts detected from the colocalization of PSDs and vesicles revealed a 52.5% or 57.2% decrease in the synapse density during QIH, compared to No-QIH or Post-QIH, respectively (Fig. 1H). This result was further confirmed by manually tracing dendrites from each group of animals (Fig. 1I, supplemental movies 1). While the density of synapses on each dendrite was significantly reduced and the distances between two neighboring synapses were larger during QIH, the volume of the remaining spines in the QIH animals was comparable to the spine volume of the awake animals (Fig. 1J). These results demonstrate a previously-unknown scale of synaptic elimination during QIH and suggest that the synaptic elimination occurs in a manner uncorrelated with spine size.

### Intact hippocampal representation and memory after QIH

The extreme structural remodeling in the hippocampus during QIH motivated us to determine if a memory and its neuronal representation acquired before QIH remain intact after QIH. We conducted multiple behavioral experiments for this purpose. First, we trained mice with the contextual fear conditioning paradigm (Fig. 2A). Animals were tested before QIH (pre) and then re-tested 5 days after recovery from QIH (post). During the post-test, both QIH mice and No-QIH control animals exhibited reliable contextual freezing in the shocked context (A) but not in the neutral context (B) with no significant difference between the pre- and post-tests (Fig. 2B). Discrimination index calculated from the freezing scores in contexts A & B indicates significant contextual discrimination in the two test sessions in both groups. Next, we tested retention of spatial memory acquired before QIH. Mice were trained to navigate to the reward location from a randomly assigned start arm on the plus maze (Fig. 2C). Once their performance met the criteria and stabilized, they entered QIH and were tested after recovery. Again, mice that had QIH after training did not show significantly different performance in the probe test compared to No-QIH controls (Fig. 2D, supplemental movie 2). We questioned whether the contextual memory remained dependent on the hippocampus after neuronal remodeling during QIH. To test the possibility that systems consolidation was facilitated during QIH, we trained mice with contextual fear conditioning and pharmacologically lesioned the hippocampus after successful memory acquisition and QIH (Fig. 2E, Fig. S2). Hippocampal lesion abolished the contextual freezing in both groups of animals, indicating that the contextual fear memory was still dependent on the hippocampal function (Fig. 2F). These results collectively show intact hippocampal memory despite the neuronal remodeling during QIH.

**Fig. 2.**
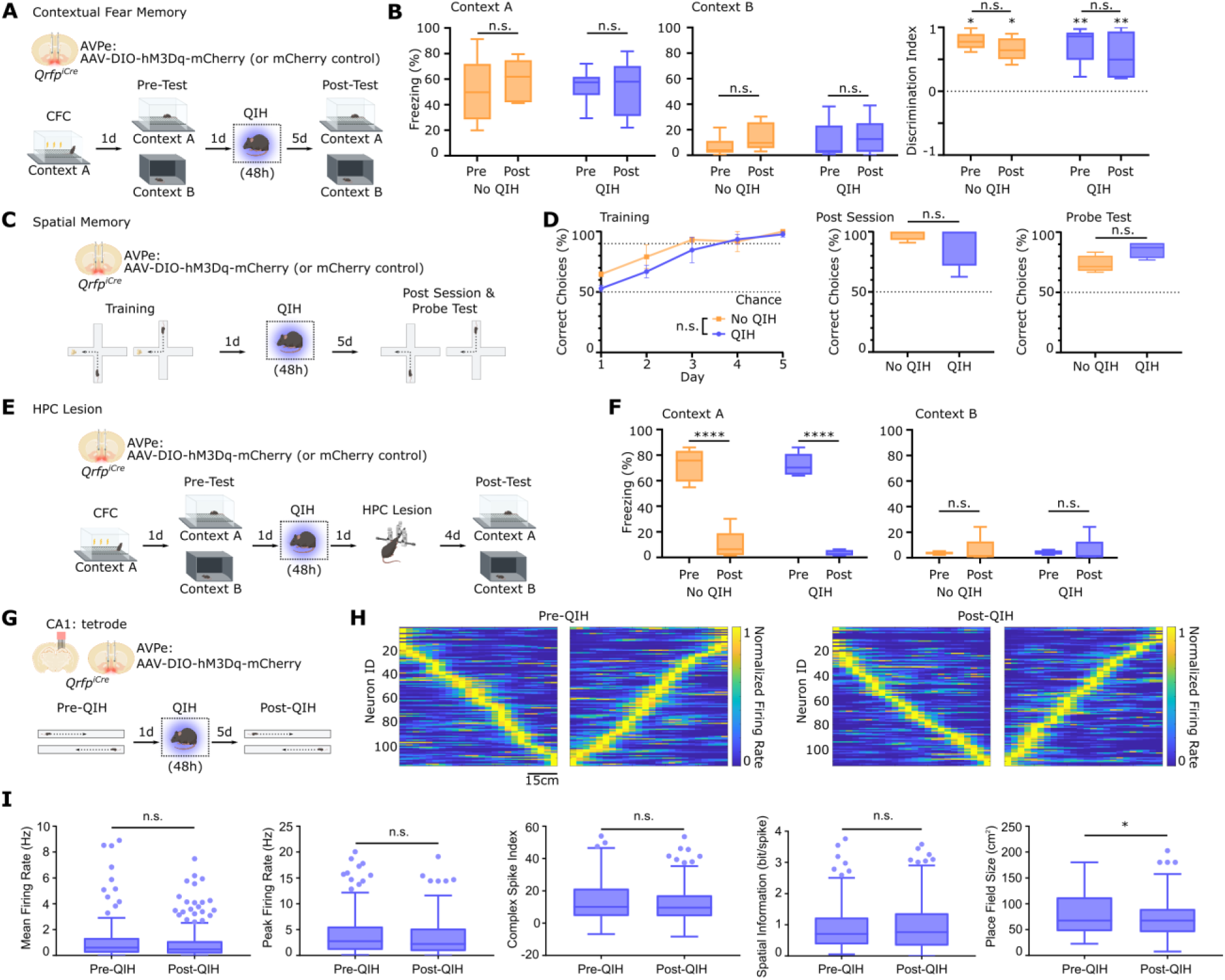
Hippocampal memories and the activity of place cells are intact after QIH. (**A**) The experimental design of the contextual fear conditioning and QIH. (**B**) Freezing scores in the shocked and neutral contexts (Context A and B, respectively) during Pre- and Post-Test in the mice with or without QIH (left) (N = 6∼10 mice for each group; Context A, Mixed Effect Model, no significant effect of the test timing or group, p = 0.2716 or 0.8599, respectively, and the multiple comparisons confirmed no difference between Pre- and Post-Test, p = 0.3089 for No QIH, p > 0.9999 for QIH; Context B, Mixed Effect Model, no significant effect of test timing or group, p = 0.0542 or 0.6515, respectively, and the multiple comparisons confirmed no difference between Pre- and Post-Test, p = 0.1976 for No QIH, p = 0.444 for QIH). (right) Discrimination index of Context A and B, one-sample Wilcoxon test for comparison with a null hypothetical value 0, p = 0.0312, 0.0312, 0.002, 0.002 for each group and session, respectively. Mixed Effect Model, a significant effect of test timing, p = 0.0424, with no significant group effect, p = 0.5261. Post-hoc tests found no significant difference between Pre- and Post-Test, p = 0.3997 or 0.1245 for No QIH or QIH, respectively. (**C**) The experimental design of the spatial memory task using the plus maze. (**D**) Performance of the spatial memory task over 5 days of training sessions (left) (N = 4 mice in each group, Mixed Effect Model, no significant effect of group, p = 0.2287). Performance during the post-session (middle) (Kolmogorov-Smirnov test, p > 0.9999), probe test (right) (Kolmogorov-Smirnov test, p = 0.1429). (**E**) The experimental design of the contextual fear conditioning, followed by QIH and a hippocampal lesion. (**F**) Freezing scores in Context A and B during Pre- and Post-Test in the mice with or without QIH (N = 5 mice for each group; Context A, Mixed Effect Model, a significant effect of lesion, p < 0.0001, and no significant effect of group, p = 0.5913. Post-hoc tests revealed significant lesion effects in both groups, p < 0.0001 for No QIH and QIH; Context B, Mixed Effect Model, no significant effect of lesion or group, p = 0.6601 or 0.9542, respectively, and the multiple comparisons confirmed no difference between Pre- and Post-Test, p = 0.9136 for No QIH, p = 0.9624 for QIH). (**G**) The experimental design of the place cell recording from the hippocampal CA1. (**H**) Heatmaps showing firing locations on the linear track, left to right laps on the left, and right to left laps on the right (n = 116 or 113 non-matched units from 3 mice for Pre-QIH or Post-QIH, respectively). Neuron IDs are sorted for each lap and session based on their peak firing locations. (**I**) Place cell properties compared between Pre- and Post-QIH. (From left) Mean firing rates, p = 0.1188; Peak firing rates, p = 0.3551; Complex spike index, p = 0.5266; Spatial information, p = 0.8761; Place field size, p = 0.0334, all Mann-Whitney tests. Box plots show median, 1st and 3rd quantiles, minimum & maximum values within 1.5 times the interquantile range (IQR) from each quantile, and the other data points outside of the range.

We then examined hippocampal representation of space before and after QIH to determine the impact of QIH on neuronal responses to external variables. Single-unit spiking activity from the dorsal CA1 was recorded when mice ran a linear track, followed by QIH induction the next day (Fig. 2G). After recovery from QIH, the animals were allowed to rerun the same track while recording spike activity. The CA1 pyramidal cells exhibited location-specific firing in both pre-and post-QIH sessions (Fig. 2H). Their average and peak firing rates did not differ between pre-and post-QIH sessions, and their complex spikes were also comparable (Fig. 2I). While the spatial information of spikes did not show a significant difference, the sizes of their place fields were significantly smaller in post-QIH than pre-QIH, indicating that spatial maps are more precise after the remodeling. These examinations did not detect any deterioration between hippocampal representations before and after the structural changes during QIH.

### Spine elimination and regrowth during QIH

The preservation of memory and its representation despite major structural changes induced by QIH suggests the reformation of a neural network equivalent to the one before QIH. To elucidate the longitudinal structural changes throughout QIH, we conducted two-photon spine imaging before, during, and after QIH induction (Fig. 3A & B). An optical path to the hippocampal CA1 of Qrfp-iCre::Thy1-YFP mice allowed dendritic imaging of the stratum radiatum stable enough to track the same dendrites over the course of 8 days (Fig. 3C & D, Fig. S3). In line with the results from EM imaging, we observed a significant elimination of dendritic spines during QIH (Fig. 3E). The fraction of the eliminated spines during the first 24 hours of QIH was significantly larger than the one during the last 24 hours before arousal (Fig. 3F). On the other hand, the normalized number of new spines which appeared during the rewarming phase of QIH was significantly larger than during the first 24 hours. These observations confirmed the extreme elimination and formation of the hippocampal dendritic spines throughout QIH, as observed in the EM dataset.

**Fig. 3.**
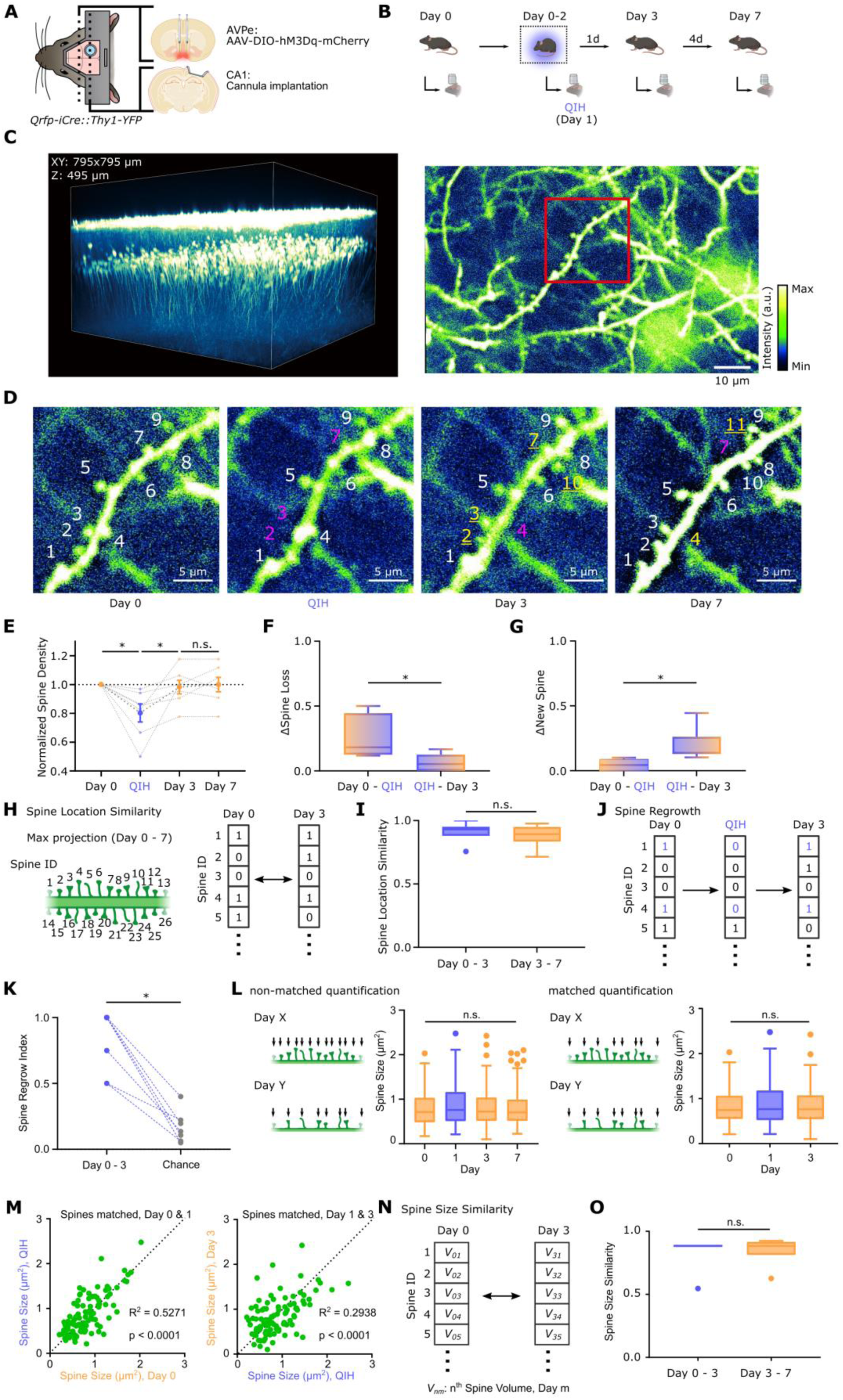
Selective elimination of dendritic spines while maintaining the sizes of the other spines during QIH. (**A**) A schematic indicating AAV injection to AVPe and cannula implantation to CA1. (**B**) The experimental design for the chronic 2p spine imaging from Day 0 to 7. After the spine imaging on Day 0, QIH was induced by injecting CNO. QIH spine imaging was conducted 24 hours after CNO injection, and mice were allowed to remain under QIH for another 24 hours and to recover for Day 3 imaging the next day in the awake state. (**C**) A 3D reconstructed image of the CA1 dendritic spines (left) and a representative image showing individual dendritic spines. (**D**) A representative image of a dendrite showing changes in its dendritic spines over the imaging sessions. The ROI corresponds to the area inside the red square in (C). Each number in the image indicates the spine ID. For Days 1-7, white-spined IDs indicate stable spines that were present in the previous imaging session and remain present in this session. Spine IDs in red indicate spines that were present in the previous session but absent in the current session. Spine IDs in yellow indicate new spines that were absent in the previous session but present in the current session. (**E**) Changes in the spine density, normalized by the number of spines on Day 0 (n = 7 dendrites from 3 mice, Friedman test, p = 0.0047, followed by post-hoc comparisons with Dunn correction, Day 0 vs QIH, p = 0.0113, QIH vs Day 3, p = 0.0214, Day 3 vs 7, p > 0.9999). (**F**) Fraction of spine loss during the first 24 hours of QIH (Day 0 – 1) and the recovery phase (Day 1 – 3) (Wilcoxon test, p = 0.0312). The number of spine loss is normalized by the total number of spines on Day 0 or 1, respectively. (**G**) Formation of new spines during the first 24 hours of QIH (Day 0 – 1) and the recovery phase (Day 1 – 3) (Wilcoxon test, p = 0.0156). The number of new spines is normalized by the total number of spines observed during the entire imaging sessions from Day 0 to 7. (**H**) A schematic indicating spine location similarity. All spines that appear at least once throughout the imaging sessions are labeled (left). In each session, a logical vector indicating the presence or absence of individual spines is assigned to each dendrite. (**I**) Cosine similarities of the spine location vectors are compared between Day 0-3 and Day 3-7 (Wilcoxon test, p = 0.4688). (**J**) Spine regrowth during QIH is defined as the presence of a spine of interest on Day 0, absence of the same spine on Day 1, and subsequent emergence on Day 3 (blue highlight). (**K**) Spine regrowth index (a ratio of regrown spines on Day 3 over lost spines on Day 1) is compared to the chance (likelihood of forming new spines at the exact locations by chance). The chance level is computed as the percentage of spines lost from Day 0 to 1 multiplied by the percentage of spines formed from Day 1 to 3 (Wilcoxon test, p = 0.0156). (**L**) The sizes of spines that are present on each day (non-matched) are compared across imaging days (left) (n = 108-129 spines, Kruskal-Wallis test, p = 0.7195). The sizes of spines that are present at Day 0, 1, and 3 (matched) are compared across imaging days (right) (n = 98 spines, Friedman test, p = 0.554). (**M**) Scatter plots comparing the spine sizes between the two imaging sessions. Pearson correlation, Day 0 vs 1 (R square = 0.5271, p < 0.0001), Day 1 vs 3 (R square = 0.2938, p < 0.0001). (**N**) Spine size similarity is the cosine similarity between two numerical vectors, indicating the volume of individual spines on specific days. (**O**) Cosine similarities of the spine size vectors are compared between Day 0-3 and Day 3-7 (Wilcoxon test, p = 0.4688). Box plots show median, 1st and 3rd quantiles, minimum & maximum values within 1.5 times the interquantile range (IQR) from each quantile, and the other data points outside of the range.

We next analyzed the spine locations on individual dendrites. More specifically, we examined the locations of newly formed dendritic spines during the rewarming phase of QIH and tested whether they formed at the same or different locations as the spine loss. We found 82.1% of spines emerged from the same locations where spines were lost during the first 24 hours of QIH (Fig. 3G). The percentage of re-emergence was significantly higher than the percentage expected based on chance alone (16.8%). We also evaluated the similarity of the locations of all dendritic spines before and after QIH. The similarity in spine locations between Day 1 and 3, with QIH in between, was 0.91 on average, which was not significantly different from the similarity between Day 3 and 7 without QIH (0.88). Finally, changes in spine sizes were examined from Day 0 to 7. The sizes of the spines identified each day (non-matched quantification) did not show significant changes across the imaging sessions, consistent with the EM observations (Fig. 3H). To better characterize the longitudinal structural changes, we extracted the spines that persisted from Day 0 to 3 (matched quantification, Fig. 3I). In contrast to the significant elimination of spines, these persistent spines do not show a significant difference in size across the imaging days. Indeed, their spine sizes during QIH were significantly correlated with and similar to the sizes before or after QIH (Fig. 3J). Congruent with this observation, the spine size similarity, including the spines eliminated and newly formed, of each dendrite between Day 0 and 3 stayed close to 1 and was not significantly different from the similarity between Day 3 and 7 (Fig. 3K). These observations using two-photon spine imaging revealed that QIH eliminated a substantial fraction of spines without altering the sizes of remaining spines, and that eliminated spines were recovered at the same dendritic locations upon arousal.

### QIH does not affect the stability of hippocampal spatial representations

We next tested whether the spine remodeling and recovery alter the hippocampal representation of experiences before and after QIH. To this end, we conducted Ca^2+^ imaging from the hippocampal CA1 using a miniature microscope. Mice were allowed to explore two distinct open fields (contexts A & B) on two consecutive days, then entered QIH the next day. After recovery, they again explored the same contexts for two consecutive days (with QIH experiment; Fig. 4A, left). We also conducted the control experiment without QIH induction between the explorations in context C & D (without QIH experiment; Fig. 4A, right). During the context explorations, Ca^2+^ dynamics in the CA1 neurons were recorded, and the cells were matched across sessions (Fig. 4B; see the methods). These cells in the hippocampal CA1 demonstrated location-specific activity and spatial information higher than their shuffled activity (Fig. 4C; see the methods).

**Fig. 4.**
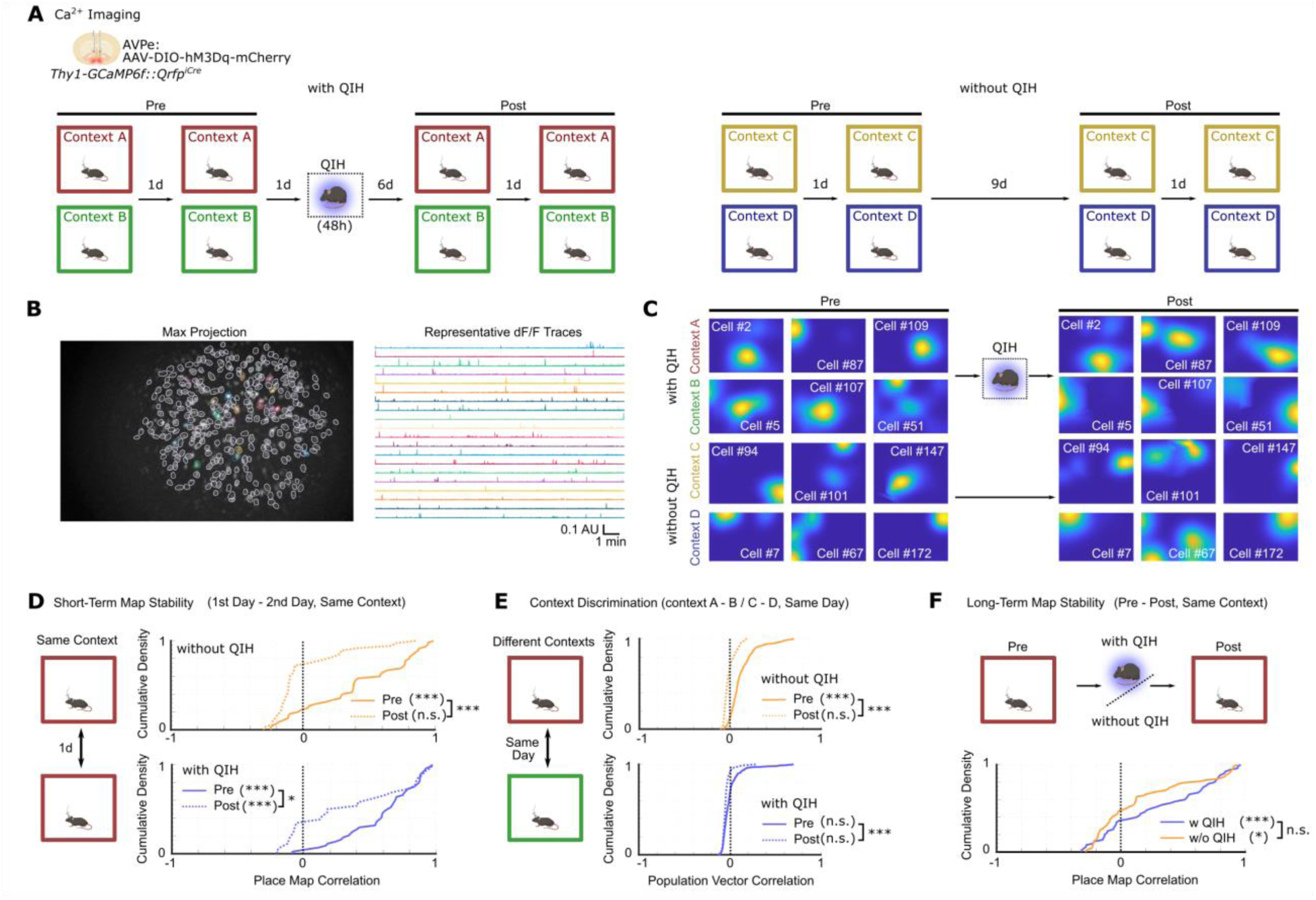
Hippocampal spatial maps remain context-specific and stable throughout QIH. (**A**) The experimental design for the chronic Ca^2+^ imaging from the hippocampal CA1 during contextual exposures before and after QIH. (**B**) A representative image indicating a maximum projection of Ca^2+^ signals (left) and their dF/F traces (right). Note that the colors of the ROIs in the max projection match those in the traces. (**C**) Heatmaps indicating representative place fields. The same sets of cells are plotted for Pre and Post-session. In the heatmaps, blue and yellow indicate the minimum and maximum Ca^2+^ activity of the cell (see the methods). (**D**) Cumulative density plots indicating the distributions of the correlations comparing spatial maps of the same context from the 1^st^ and 2^nd^ day during Pre- or Post-sessions (without QIH (top), n = 45 (Pre) and 36 (Post) place cells from N = 3 mice, one-tailed (right-tail) Wilcoxon signed rank test testing whether median > 0, p < 0.0001 for Pre, p = 0.852 for Post, Mann-Whitney U test for group difference, p < 0.0001; with QIH (bottom), n = 44 (Pre) and 26 (Post) place cells from N = 3 mice, one-tailed Wilcoxon signed rank test, p < 0.0001 for Pre, p < 0.0001 for Post, Mann-Whitney U test for group difference, p = 0.035). (**E**) Cumulative density plots indicating the distributions of the population vector (PV) correlations comparing spatial maps of the different contexts on the same day during Pre- or Post-sessions (without QIH (top), n = 440 (Pre) or 440 (Post) bins, Wilcoxon signed rank test (right-tail), p < 0.0001 for Pre, p > 0.999 for Post, Mann-Whitney U test for group difference, p < 0.0001; with QIH (bottom), n = 494 (Pre) or 494 (Post) bins, Wilcoxon signed rank test (right-tail), p = > 0.999 for Pre, p = > 0.999 for Post, Mann-Whitney U test for group difference, p < 0.0001). (**F**) Cumulative density plots indicating the distributions of correlations comparing spatial maps of the same context between Pre- and Post-sessions (n = 68 (without QIH) or 50 (with QIH) place cells from N = 3 mice, Wilcoxon signed rank test (right-tail), p = 0.0118 for without QIH, p < 0.0001 for with QIH, Mann-Whitney U test for group difference, p = 0.0908). Box plots show median, 1st and 3rd quantiles, minimum & maximum values within 1.5 times the interquantile range (IQR) from each quantile, and the other data points outside of the range.

These place cells showed significant similarity between the two spatial maps from sessions 24 hours apart in the same context, indicating short-term map stability during Pre-QIH sessions with or without QIH (Fig. 4D). Over 9 days of the interval between Pre- and Post-sessions, however, the stability of place cells decreased in both experiments. Notably, place cells remained stable after QIH but became unstable after the interval without QIH, suggesting that QIH led to greater stabilization of place cells. We next examined the context-specificity of place cells (Fig. 4E).

Because there was a smaller overlap of place cells between explorations in different contexts, we used population vectors (PVs) that included cells that were inactive in one of the two imaging sessions (*44*). The PV correlations between two different contexts in Post-sessions were significantly smaller than in Pre-sessions in both experiments with and without QIH, indicating better context discrimination after the interval, regardless of QIH. Finally, long-term map stability between Pre- and Post-sessions was examined (Fig. 4F). Because hippocampal spatial maps decorrelate over time even without QIH (representational drift; (*21*, *22*)), we directly compared map similarity between with and without QIH. In both experiments, place maps from two imaging sessions, 10 days apart, demonstrated significant similarity. There was a slight trend toward greater similarity in the QIH experiment, but no significant difference was observed between with and without QIH, indicating that QIH did not decrease map stability more than the degree produced by passage of time. Overall, these longitudinal observations of Ca^2+^ dynamics showed that QIH did not deteriorate the integrity of the hippocampal representations of space.

### Clustered engram synapses are spared during QIH

Our results so far have characterized the intense remodeling of the hippocampal neuronal network during QIH and yet indicate remarkably intact retention of dendritic structures, hippocampal representations, and memory. To compare structural changes that lead to memory impairment with those that do not, we conducted a series of experiments using long-term anesthesia and pharmacological blockade of actin polymerization (ANE), rather than QIH, as a condition that resembles behavioral inactivity and synaptic disruption (*47*, *48*). Similar to QIH, ANE produced a significant reduction in firing rate (Fig. 5A), oscillation amplitude (Fig. 5B), and synapse density in the hippocampus (Fig. 5C). In contrast to QIH, however, mice exhibited a significant impairment in contextual freezing during the post-test after recovery (Fig. 5D & E).

**Fig. 5.**
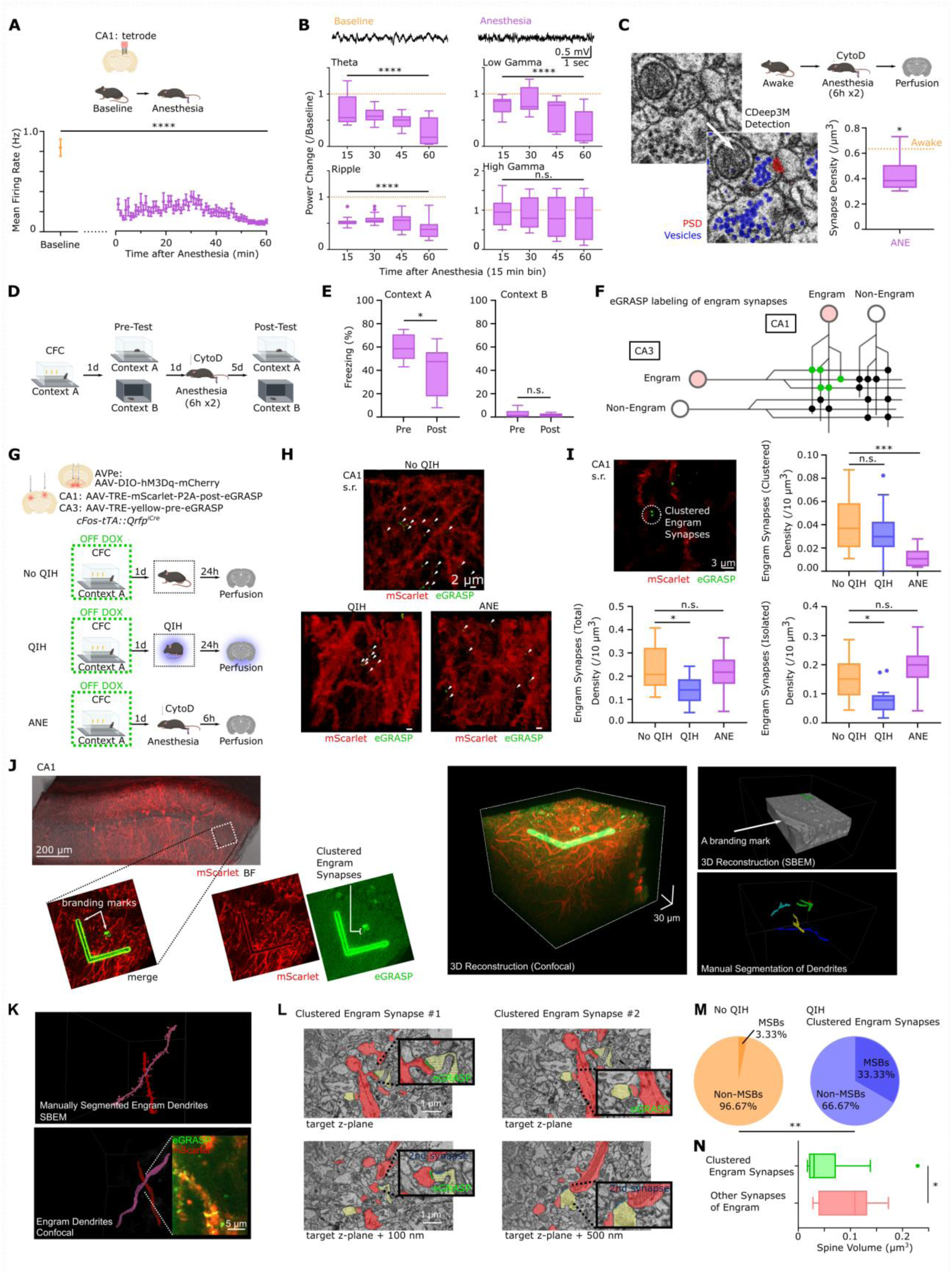
Clustered engram synapses are preferentially spared during QIH. (**A**) The firing rate changes of the hippocampal neurons (the CA1 pyramidal cells) during anesthesia (n = 228 units from 3 mice, Friedman test, p < 0.0001). Plots show the firing rate values averaged over 15 minutes of the resting session (Baseline) or over 1-minute bins after mice are anesthetized. Error bars indicate the standard error of the means. (**B**) (Top) A representative LFP trace from a Baseline (left) or anesthesia (right) session. (Bottom) Changes in the oscillation power after anesthesia (n = 23 recording channels in the hippocampus from 3 mice). Theta (4-12 Hz, p < 0.0001), low gamma (30-50 Hz, p < 0.0001), high gamma (65-85 Hz, p = 0.2725), and ripple (100-130 Hz, p < 0.0001) ranges were examined (Friedman test). The mean power in the baseline session normalizes power values. (**C**) Synaptic density within a ROI (850 um^3^) in ANE mouse brains (6 hours of anesthesia twice in 2 successive days, followed by Cytochalasin D (CytoD) intra-hippocampal infusion, see the methods for detailed procedure) compared with No-QIH animals (right bottom) (n = 29 ROIs from 3 mice, One-Way ANOVA followed by post-hoc multiple comparisons with Dunnett’s correction, p = 0.0359 for No-QIH vs ANE). Synapses are identified using CDeep3M detection of PSD and pre-synaptic vesicles(left). (**D**) The experimental design of the contextual fear conditioning and ANE. (**E**) Freezing scores in the shocked and neutral contexts (Context A and B, respectively) during Pre- and Post-Test in the mice with or without ANE (left) (N = 8 mice; Context A, Wilcoxon test, p = 0.0234; Context B, Wilcoxon test, p = 0.8594). (**F**) A schematic indicating eGRASP engram synapse labeling. Pre-and Post-eGRASP are expressed in CA3 and the contralateral CA1, respectively. (**G**) A diagram indicating the surgical design (top) and the three groups of mice for eGRASP labeling at different states (bottom). See methods for detailed procedure. (**H**) 3D reconstructed confocal images showing the dendrites of engram cells (red) and eGRASP puncta along the dendrites (green dots, pointed by green arrows). S.R., the stratum radiatum. (**I**) A representative confocal image showing the dendrites of engram cells (red), eGRASP puncta (green), and clustered engram synapses (two eGRASP puncta within 5 μm distance, white blanket) (top left). (bottom left) The density of all engram synapses in the three groups (n = 12∼17 ROIs from 3 mice each group, Kruskal-Wallis test, p = 0.0157, followed by multiple comparison tests with p value corrected by Dunn’s method, NoQIH vs QIH, p = 0.029, NoQIH vs ANE, p > 0.9999). (top right) The density of clustered engram synapses in the three groups (Kruskal-Wallis test, p < 0.0001, followed by multiple comparison tests, NoQIH vs QIH, p > 0.9999, NoQIH vs ANE, p = 0.0002). (bottom right) The density of isolated engram synapses in the three groups (Kruskal-Wallis test, p = 0.0005, followed by multiple comparison tests, NoQIH vs QIH, p = 0.0242, NoQIH vs ANE, p = 0.5823). (**J**) A schematic indicating CLEM (correlative light and electron microscopy) to identify and observe clustered engram synapses with fluorescent labels. The clustered eGRASP puncta are first identified via confocal microscopy. Two distinct branding marks are created in the close vicinity of the target eGRASP signals (left). The confocal and electron microscopic images are 3D-reconstructed to identify the same synaptic contacts in EM images based on the locations of the branding marks and the spatial arrangements of dendrites, nuclei, and blood vessels (right). (**K**) Representative engram dendrites were manually 3D reconstructed from SBEM images (top) and matched with dendrites with eGRASP synapses from confocal images (bottom). (**L**) Example SBEM images indicating eGRASP-positive synapses (green highlights) belonging to a cluster, and their pre-synaptic partners (yellow highlights). Note that their pre-synaptic partners are multi-synaptic boutons (another synapse on the same bouton is indicated in blue highlight). (**M**) Pie plots indicating the proportion of multi-synaptic boutons and single synaptic boutons from randomly selected spines in No QIH mice (left) (n = 60 from N = 3 mice), and the proportion of multi-synaptic boutons and single synaptic boutons from clustered eGRASP-positive synapses in QIH mice (right) (n = 12 from N = 2 mice, Fisher’s exact test, p = 0.0059). (**K**) Spine volumes compared between clustered engram synapses and the other synapses on the same engram dendrites from QIH animals (Mann-Whitney test, p = 0.0332). Box plots show median, 1st and 3rd quantiles, minimum & maximum values within 1.5 times the interquantile range (IQR) from each quantile, and the other data points outside of the range.

These results highlight distinct effects of hippocampal changes on memory between QIH and ANE and suggest that QIH is a controlled hypometabolic and hypothermic state that maintains the structural integrity of memory retention.

To elucidate the structural underpinning of memory, which should be spared under QIH but compromised under ANE, we labeled synaptic connections between c-Fos-tagged engram neurons using eGRASP (green fluorescent protein reconstitution across synaptic partners) (Fig. 5F, (*47*, *48*)). Engram neurons in CA1 and CA3 of the hippocampus during contextual fear conditioning were tagged, allowing for expression of Post-eGRASP and Pre-eGRASP, respectively. Animals then entered QIH or ANE on the subsequent day (Fig. 5G). The density of eGRASP puncta along mScarlet-labeled engram dendrites exhibited a significant decrease in QIH animals compared to No-QIH controls, indicating a substantial elimination of engram-engram synapses during QIH (Fig. 5H & I, bottom left). The eGRASP puncta in the ANE animals showed only a trend of decrease. These results showed that the densities of engram-engram synapses alone do not explain memory retention after QIH or memory impairment after ANE.

Recent studies reported spatially clustered engram-engram synapses in the hippocampus (*48*, *49*). In line with these studies, we also observed clustering of eGRASP puncta in the non-QIH control animals (Fig. 5H & I, Fig. S4). Notably, we found the density of the clustered eGRASP puncta in the QIH animals was comparable to that in the control animals, while non-clustered eGRASP puncta in the QIH mice were significantly decreased (Fig. 5I, top right). In contrast, the clustered eGRASP puncta in the ANE mice were significantly reduced, while non-clustered eGRASP puncta did not show a significant change (Fig. 5I, bottom right). This finding indicates that clustered engram-engram synapses are preferentially protected during QIH, while such selective protection does not occur during ANE. To further characterize the network architecture of the clustered engram-engram synapses during QIH, we specifically examined their ultrastructure and post-synaptic axons using SBEM and CLEM (correlative light and electron microscopy) (Fig. 5J). The clustered engram-engram synapses in CA1 were identified by confocal microscopy and tagged with laser-branding marks for identification under EM. Three-dimensional reconstruction of engram dendrites from confocal images and corresponding dendrites from SBEM images identified clustered engram synapses from the EM images (Fig. 5K). We found 33.33% of eGRASP-positive clustered spines make synaptic contacts with multi-synaptic boutons (MSBs; (*50*)) (Fig. 5L & M). The percentage of synaptic contacts with MSBs observed in the clustered engram synapses during QIH was significantly higher than the percentage observed in randomly selected synapses from No QIH animals (3.33%) (Fig. 5M).

Importantly, MSBs making synaptic contact with one of the clustered engram synapses have synaptic contacts with spines from other dendrites, suggesting that the clustered engram-engram synapses facilitate coordination of population activity across neurons, rather than enhancement of post-synaptic response from a single pre-synaptic axon. Moreover, we found that the spine volumes of the clustered engram-engram synapses were not significantly larger, rather significantly smaller, than other spines on the same engram dendrites from QIH animals, further strengthening the view that larger spines are not necessarily preserved during QIH (Fig. 5N). These observations uncovered a distinct neuronal architecture that was spared during the massive spine elimination during QIH.

## Discussion

The structural underpinning of memory has been one of the most fundamental topics in neuroscience for decades. While ample experimental evidence has accumulated to show that experience-dependent and activity-driven synaptic plasticity is indispensable for successful memory encoding, there is high turnover of dendritic spines and continuous drifting of the neuronal representations of experience (*19–24*). This may suggest that long-lasting changes introduced during memory encoding do not need to persist throughout memory retention (*52*, *53*). However, an implicit assumption in current models of memory storage is that there are enduring structural changes which underlie long-term memory storage. If that is the case, what could be the structural underpinning that enables long-term storage of memory in the ever-changing neuronal network?

In the current study, we used artificial hibernation in mice to induce extreme downscaling of the neuronal activity and dendritic structures in the hippocampus. During this hypothermic and hypometabolic state, hippocampal neurons show a ∼70% reduction in firing rate and the elimination of more than 50% of synapses. A straightforward prediction from this phenomenon, based on the widely accepted idea of synaptic memory trace, would be a significant memory impairment after arousal. Contrary to this prediction, our study reveals intact hippocampal memory despite drastic changes in the neuronal network. We also demonstrate that spatial representations in the hippocampus are intact after QIH. Surprisingly, spine changes during QIH differ from global synaptic scaling during sleep that involves proportional downscaling over all spines homogeneously (*54–60*). In the case of QIH, while many spines are eliminated, the remaining spines maintain their size throughout QIH. These protected spines have larger PSDs, but spine sizes are comparable to the average during awake, suggesting their unique ultrastructural and molecular signatures that we did not report in the current study. Another unexpected finding is the structural recovery of the dendritic spines upon arousal. Most of the eliminated spines regrow from the same dendritic locations, suggesting the reconstruction of the neuronal network congruent to the one before QIH. In line with this view, we also found that spatial representations in the hippocampus are still context-specific and stable after QIH. It needs to be emphasized that, however, the current study did not determine whether presynaptic counterparts of regrown spines and their synaptic efficacies were also the same upon the spine regrowth. Future studies are needed to fully elucidate the changes in neuronal connections throughout QIH.

These unique forms of spine remodeling during QIH raise two hypotheses regarding the mechanism of memory retention in the ever-changing neuronal network. The first hypothesis is that a core (and sparse) memory trace underlies memory retention. This core memory trace does not require spines that are enlarged at memory encoding to remain enlarged throughout memory retention. Under this framework, spines that belong to the core memory trace and survive QIH are not necessarily larger but are selectively protected during the massive spine elimination. On the other hand, the second hypothesis does not assume that the structural underpinning of memory is intact but instead assumes a latent capability to reconstruct a consistent network after alteration and perturbation, as observed in the two-photon experiment of the current study. In this scenario, one can predict intact memory even when all spines are eliminated, if the latent capability is not compromised and can later reconstruct the functional network for all stored memories.

These two hypotheses are not mutually exclusive, and it is possible that the core memory trace contributes to the larger reconstruction of the original network upon arousal from QIH. However, past studies of awake animals support the view of the core memory trace. First, long-term spine imaging or Ca^2+^ imaging revealed highly decorrelated dendritic spines and hippocampal representations after days, suggesting that the network does not need to be identical or reconstructed for the sake of memory. Second, the hippocampus is known to be an automatic encoder of episodic memory (*54*, *55*), making this structure profoundly plastic. If the same pattern of synaptic connections is the key to successful memory retention, the hippocampus becomes highly vulnerable to interference caused by continuously encoded episodic memories. Sparse core memory traces could be more easily embedded into the automatic encoding system with less interference among multiple memory traces.

The current study uncovered the spatially clustered engram-engram synapses and their associated architecture as a core memory trace. Past studies reported clustered synaptic potentiation in hippocampal dendrites and proposed its relevance to memory (*56–62*). This hetero-synaptic principle facilitates supra-linear integration of synaptic inputs and detection of specific input patterns (*63–66*). Moreover, a study found that high spine turnover promotes the formation of clustered spines, suggesting that a highly plastic hippocampal network is beneficial, rather than destructive, for the use of this particular structural motif as the core memory trace (*58*). Despite these insightful findings on clustered spines, their relevance to memory remains observational, due to experimental difficulties in de-clustering target spines without altering other variables (*58*). In the current study, by using artificial hibernation, we took a distinct approach to downscale the neuronal network to the structural essentials without impairing memory retention. Even after more than half of synapses are eliminated, the clustered engram synapses are spared and maintain their synaptic architecture—spine sizes and connections to MSBs. In contrast, animals in a behaviorally inactive state with synaptic perturbation exhibit memory impairment and have fewer clustered engram synapses in the hippocampus. Our findings provide strong evidence that clustered engram synapses are sufficient for memory retention, whereas non-clustered engram synapses are dispensable. Along with other results in the current study, this conclusion suggests that the biological trace of memory retention is not necessarily associated with larger spines, is even sparser than our current model, and requires a particular hetero-synaptic motif, resulting in a scattered, broader neuronal network for memory storage. A deeper understanding of the mechanism by which hibernation protects the core memory trace and maintains the network integrity would reveal more comprehensive principles of memory.

## Acknowledgments

We would like to thank Thomas J. McHugh at RIKEN, Joshua P. Johansen at RIKEN, Yoko Yazaki-Sugiyama at OIST, Inna Slutsky at Tel Aviv University, and David Dupret at University of Oxford for comments and discussions on an earlier version of the manuscript. We also appreciate all the members of the Memory Research Unit and the supporting sections in OIST. We especially thank K. Nakamura for daily assistance with our research activities, the Animal Resource Section for the maintenance of our mouse colony, M. Kuroda and C. Swart in the Engineering Section for 3D printing Microdrive parts and construction of sample holders for EM, M. Hall in the Scientific Imaging Section for assistance and maintenance of EM machines, and the members of Kubota and Ohno labs at NIPS for technical advice for the CLEM experiment. We are grateful to Yuki Watakabe, Maki Watanabe, Sachiko Furukawa, Sanae Hara, and Chiemi Hyodo for their help with animal care in NIPS. We also thank Arthur J. Y. Huang at RIKEN for DNA plasmids, AAV vectors, and technical advice on AAV production. Some illustrations in figures are created with BioRender.com. This work is supported by OIST institutional research budget (KZT), JSPS KAKENHI Grant-in-Aid for Scientific Research (B) (Grant Number: 21H02585, KZT), Grant-in-Aid for Transformative Research Areas (A) (Grant Number: 23H04944 & 23H04939, KZT), and CREST (Grant Number: JPMJCR24T4, KZT), NIPS and Tokyo university of science institutional research budget (TN, RE, TT), JSPS KAKENHI Grant-in-Aid for Scientific Research (S) (Grant Number: 20H05669, TN, RE, TT), Grant-in-Aid for Transformative Research Areas (B) (Grant Number: 20H05769, RE), Fund for the Promotion of Joint International Research (Grant Number: 22K21353, TN), Grant-in-Aid for Transformative Research Areas (A) (Grant Number: 22H04926, TN), Grant-in-Aid for Scientific Research (B) (Grant Number: 23H04943, RE), Grant-in-Aid for Scientific Research (A) (Grant Number: 25H01025, TN), and Grant-in-Aid for Early-Career Scientists (Grant Number: 23K14294, 25K18950, TT), AMED Brain/MINDS 2.0 (Grant Numbers: JP24wm0625105, TN; JP24wm0625211, RE, TT; and JP24wm0625109, TT), the “Advanced Bioimaging Support” project, the “NINS Program of Promoting Research by Networking among Institutions” (Grant Number: 01412303), the Nakatani Foundation (Grant Number: 2025S206, TT), the “Joint Research Program of the Exploratory Research Center on Life and Living Systems (ExCELLS)” (Grant Numbers: 21-205, 22EXC202, 23EXC337, 23EXC601, 24EXC301, 25EXC305, 25EXC501), Frontier Photonic Sciences Project of NINS (Grant Number: 01212505), and Cooperative Study Programs of National Institute of Physiological Sciences (KZT).

## Author contributions

Conceptualization: KZT

Methodology: YJL, ATN, KT, TT, DM, HA, MCC, MH, MU, AM, YM, RE, AH, SL, NO, YK, TS, KZT

Investigation: YJL, ATN, KT, TT, DM, HA, MCC, YM, SL, NO, YK, KZT

Formal Analysis: YJL, ATN, KT, TT, DM, MU, YM, SL, NO, YK, KZT

Visualization: YJL, ATN, KT, TT, DM, YM, SL, NO, YK, KZT

Funding acquisition: KZT

Project administration: KZT

Resources: YJL, ATN, KT, TT, DM, HA, MCC, MH, MU, AM, YM, TN, RE, AH, KS, SL, NO, YK, TS, KZT

Supervision: KZT

Writing – original draft: KZT

Writing – review & editing: YJL, ATN, KT, TT, DM, HA, MCC, MH, MU, AM, YM, TN, RE, AH, KS, SL, MO, YK, TS, KZT

## Competing interests

Authors declare that they have no competing interests.

## Supplementary Materials

Materials and Methods

Figs. S1 to S4

References (*67–70*)

Movies S1 to S2

## Notes

### Competing Interest Statement

The authors have declared no competing interest.

