## Supplemental Information for "Artificial hibernation uncovers distinct synaptic engram architecture for memory retention"

**The PDF file includes:**

Materials and Methods
Figs. S1 to S4
References

**Other Supplementary Materials for this manuscript include the following:**

Movies S1 to S2

### Materials and Methods

Subject

Qrfp-iCre mice were generated and maintained on a C57BL/6J background by backcrossing for at least 10 generations, as previously described (*38*). c-Fos-tTA mice (JAX 018306) or Thy1-GCaMP6f (JAX 024276) mice, or Thy1-YFP (JAX 003782) were crossed with Qrfp-iCre mice to obtain F1 hybrid animals carrying both transgenes. Both males and females were randomly assigned to each of the experimental groups in the study. Qrfp-iCre::c-Fos-tTA mice, including their breeders, were provided water and doxycycline-containing food (40 mg/kg, Bioserv) *ad libitum* and maintained in a temperature- and humidity-controlled room with a 12 h light/dark cycle (lights on from 07:00 A.M. to 7:00 P.M.) under the standards of AAALAC accreditation. All experimental protocols were approved by the OIST Animal Care and Use Committee or the Institutional Animal Care and Use Committee of the National Institute of Natural Sciences.

Surgery

AAVs were either purchased from Addgene or produced using AAVpro Purification Kit (Takara, Japan). Titers of the vectors were as follows: AAV.hsyn.DIO.hM3D(Gq).mCherry, 2.30 x $10^{13}$; AAV.hSyn.DIO.mCherry, 1.70 x $10^{13}$; AAV.TRE.mScarlet.P2A.post.eGRASP, 1.22 x $10^{12}$; AAV.TRE.yellow.pre.eGRASP, 1.17 x $10^{12}$ (copies/mL).

All animals were single-housed after surgery. Mice were anesthetized with either isoflurane (1.5% for induction and 1.0% for maintenance) or i.p. injection of MMB (a mixture of midazolam, medetomidine, and butorphanol tartrate at concentrations of 4.0 mg/kg, 0.3 mg/kg, and 5.0 mg/kg, respectively) and placed on the stereotaxic apparatus (Kopf Instruments). For infusion to AVPe (AP, 0.38 mm; ML, $\pm$0.3 mm; DV, -5.1 mm), 300 µL of AAV-DIO-hM3Dq-mCherry or AAV-DIO-mCherry was injected. For infusion to CA1 (AP, -2.0 mm; ML, 1.5 mm; DV, -1.3 mm), 500 µL of AAV-TRE-mScarlet-p2A-post-eGRASP was injected. For infusion to CA3 (AP, -1.9 mm; ML, -2.35 mm; DV, -2.45 mm), 500 µL of AAV-TRE-pre-eGRASP was injected. More specifically, each AAV was infused through a small craniotomy above the site at a rate of 100 µL per minute with a 33G blunt needle (NF33BL, NanoFil) and a micropump syringe (World Precision Instrument). After infusion, the 33G needle stayed at the infusion site for 5 minutes to allow diffusion of the vector.

The microdrives for single-unit and LFP (Local Field Potential) recording were constructed as previously described (*67*). It had eight independently adjustable tetrodes (14 nm diameter, nichrome wire, gold-plated to an impedance of 200-300 kΩ) and another adjustable tetrode for reference. The Microdrive was implanted to target the dorsal CA1 (AP, -2.0 mm; ML, 1.5 mm; DV, -1.3 mm). Five stainless head screws are used to anchor the Microdrive on the skull. One of the stainless screws was used as an electrical ground for extracellular recording. The Microdrive was cemented using two layers of dental cement (Refine Bright acrylic resin, Yamahachi Dental, Japan; Super Bond, Sun Medical Co., Ltd, Japan). After recovery, the depth of the tetrodes was slowly and manually adjusted until stable recordings of CA1 units and reliable detection of sharp wave ripples were achieved. During the adjustments, mice stayed in a highly familiar small plastic bucket (20 cm diameter) located in the recording room.

For one-photon $Ca^{2+}$ imaging, GRIN lens (ProView Integrated Lens, 1.0 mm diameter, 4.0 mm lens length, Inscopix, USA) was implanted at the dorsal CA1 of the hippocampus (AP, -2.0 mm; ML, 1.5 mm; DV, 1.6 mm). Cortical tissue above the target corpus callosum was aspirated using 26G and 27G needle to secure the lens path.

For in vivo two-photon imaging, a craniectomy followed by cortical removal and cannula implantation was performed. Mice were anesthetized with isoflurane (3.0% for induction and 1.5% for maintenance), and body temperature was maintained using a disposable heating pad. To reduce intracranial pressure and loosen the dura, i.p. injection of GLYCEOL (TAIYO Pharma) was administered 15 minutes before craniectomy. After removing the hair and skin to expose the skull, an approximately 4 mm diameter craniectomy was performed on the CA1 (AP, -2.0 mm; ML, 2.0 mm). The exposed dura mater and the underlying cerebral cortex were removed using fine tweezers and a suction pump. Finally, a custom-made metal cannula with a glass coverslip (2.7 mm in diameter; No. 3, Matsunami) was implanted into the brain hole to the target depth (DV, -1.0 mm) and secured to the skull with a cyanoacrylate adhesive or dental cement (Ionosit baseliner, DMG). Following the surgery, the mice were allowed to recover for at least 3 weeks before the start of the experiment.

QIH induction

Before QIH induction, mice were handled by an experimenter for at least 5 days. Two to three weeks after surgery, 0.1 mL/g of saline was injected intraperitoneally (i.p.) once daily for 4 days to habituate the i.p. injection. The day before the QIH injection, the mice were shaved on the back to measure surface body temperature.

On the day of QIH induction, mice were allowed to habituate in the hibernation chamber (HC-10, Shin Factory) for 1 hour. The chamber is air-circulated, set at 20 °C, and under constant darkness. Within the hibernation chamber, each animal was placed in a black plastic box (9 x 18 x 20 cm, no ceiling) with bedding materials, water gel, and food pellets provided. Their body temperature was recorded using the infrared thermal camera (FLIR C5) attached 50 cm above the box. The animal's temperature and location were identified and tracked using custom-made software (HIBER; see https://github.com/oist/Tanaka_K_Lab_public for details).

After habituation, CNO (1 mg/kg) was i.p. injected to activate hM3Dq-expressing Q-neurons in AVPe. Mice were allowed to stay in the hibernation chamber for 48 hours, during which their temperatures and locations were recorded at 1 Hz using the IR camera and HIBER. After 48 hours, the mice were returned to their home cage in the vivarium and allowed to recover without disturbance.

In in vivo spine imaging, body temperature was recorded from the shaved skin area using an infrared thermal camera (Pl450i, Optris). CNO (1 mg/kg) was i.p. injected to activate hM3Dq expressed in Q-neurons within the AVPe. QIH induction and observation were performed in a head-fix condition under a two-photon microscopy (A1RMP+, Nikon) in a dark room maintained at approximately 21 °C. After QIH induction, the mice were kept under the same conditions for 120 min and then returned to their home cages in the vivarium for undisturbed recovery.

Thermal Imaging Acquisition and Analysis System (HIBER, Heat Information and Behavioral Estimation Recorder)

Data were acquired using HIBER, a custom C++ application developed for real-time processing of multiple thermal cameras. The software relies on the Qt Framework, the QCustomPlot library, and OpenCV for video stream handling and image processing. Because the FLIR C-5 cameras do not support software readout of the temperature sensor, HIBER extracts temperature data in real time from video frames. Frames are streamed to memory using the DirectShow interface. A predefined region of interest (ROI) is applied to remove watermarks and dashboard artifacts burned onto the image. Each ROI can be subdivided into multiple subregions, enabling independent analysis of distinct experimental arenas. Raw pixel values from thermal cameras are converted to calibrated temperatures (°C) by linear mapping based on the user-selected temperature range. HIBER then performs image processing to isolate the animal shape using background subtraction and to compute metrics, including the minimum, mean, maximum, and standard deviation of temperature. Object tracking is achieved by identifying the centroid coordinates determined through thresholding and contour-based image analysis. The software is available in the public repository (https://github.com/oist/Tanaka_K_Lab_public).

Tetrode recording

All tetrode data were acquired using a 32-channel Digital Lynx SX acquisition system (Neuralynx). LFP was filtered between 2 and 9000 Hz. Spike waveforms were filtered between 0.6-6 kHz, and those above a peak threshold of 50 μV were time-stamped and digitized at 32,556 Hz. For place cell recording on a linear track, light-emitting diodes (red and green) on the recording headstage were video-tracked to obtain the animal's position and head direction with a sampling rate of 30 Hz.

After all recording sessions, mice were anesthetized for electrolytic lesions to mark tetrode locations and then transcardially perfused. Following 24 hours of post-fixation with 4% PFA (paraformaldehyde), brains were sectioned (50 μm thickness) to confirm tetrode locations in the hippocampal CA1.

Spike sorting

Spikes were manually sorted using SpikeSort3D software (Neuralynx) as previously described (*67*). The waveform variables (peak amplitudes and their energies) of all spikes recorded from a combination of three channels were projected to 3D space, and putative units were manually clustered. This procedure was repeated for all channel combinations. Clusters with > 0.5 % spikes having an inter-spike-interval of < 2 msec, a total number of spikes < 50, or an isolation distance < 10 were excluded.

Clustered units were matched between sessions (before and 0-2 hrs after QIH induction) as previously described (*67*). Briefly, units that passed our selection criteria between sessions were considered the same only when a slight boundary adjustment was sufficient to match them.

Processing $Ca^{2+}$ imaging data

Calcium imaging videos were preprocessed using Inscopix Data Processing Software for spatial filtering, motion correction, and ΔF/F computation. Neurons were identified by manually delineating regions of interest (ROIs) exhibiting clear calcium increase from baseline. To track cells across sessions, we implemented a MATLAB-based cell-matching algorithm. The median projection image from the first session served as the fixed template, and subsequent sessions were aligned to it via affine transformations estimated using SURF feature matching and cross-correlation optimization, selecting the transformation leading to the highest correlation with the template. The resulting transformation was applied to all ROIs, projecting them into the template coordinates. ROIs were considered to represent the same neuron if their containment ratio (intersection area divided by the smaller ROI area) exceeded 0.7. Cells detected in at least two sessions were assigned unique identifiers; others were classified as singletons. Neural activity was temporally aligned with behavioral data via TTL signals between acquisition systems.

Place fields

The mice's trajectories were corrected for artifacts caused by transient tracking errors, then smoothed with a Gaussian kernel of 0.05 SD width. The trajectories were then assigned to left or right laps based on their direction of motion. Firing rate maps were obtained by assigning each cell's spikes to 3 cm spatial bins when the animal moved faster than 2 cm per second, then dividing by the bin's occupancy time. The firing rate maps were then ordered by peak firing rate location.

Pyramidal cells with a peak firing rate > 3 Hz and spatial information > 0.3 bits/spike were classified as place cells. Mean firing rates were the average firing rates of each unit when it exceeded the speed threshold (2 cm per second). Peak firing rates were the highest values in the firing rate maps. The place field size was defined as the number of spatial bins that exceeded 20% of its peak firing rate. Spatial information and Complex Spike Index (CSI) were computed as previously described (*68*). Interneurons (cells having mean firing rates > 10 Hz and place field size > 1/3 of the entire area) were excluded from the analysis.

For calcium imaging data, pseudo–firing rate maps were computed as the mean ΔF/F activity associated with calcium transients within each spatial bin. For this, the arena was divided into 2 × 2 cm spatial bins. The animal's head position was estimated using a background-subtraction and ellipse-tracking algorithm incorporating a constant-velocity Kalman filter. Periods of immobility (velocity $< 2$ cm $s^{-1}$) were excluded from the analysis. Fluorescence traces (ΔF/F) were smoothed with a Gaussian kernel with a window length of 1s and detrended by subtracting the local baseline defined as the 20th percentile within a 5 s sliding window. For each neuron, the mean and standard deviation of the detrended signal were computed. Frames in which ΔF/F exceeded the mean by more than two standard deviations were marked as candidate calcium transients. Events shorter than 300 ms were discarded, and events separated by < 500 ms were merged into single transients. Imaging frames (20 Hz) were synchronized with behavioral video frames (30 Hz) using TTL pulses transmitted from the Inscopix nVista acquisition system to the Neuralynx Digital Lynx recorder.

Spatial tuning was quantified using spatial information (SI). Statistical significance was assessed by generating a null distribution of SI values through 1000 shuffle iterations in which each neuron's calcium-transients were randomly placed in the timeline while preserving its temporal structure. Neurons whose observed SI exceeded the 95th percentile of the shuffled distribution were classified as place cells. Spatial pseudo–firing rate maps were constructed by accumulating calcium-transient amplitudes within each 2 × 2 cm spatial bin and normalizing by occupancy time. Both occupancy and amplitude maps were smoothed with a Gaussian kernel ($\sigma = 1$ bin) before division to generate the final pseudo-firing rate maps.

Spatial stability across sessions or contexts was quantified using place-field (PF) correlation, computed only for neurons classified as place cells in both conditions. The Pearson correlation coefficient (r) between their two-dimensional rate maps from the compared sessions was used to measure place-field stability. Population-level stability was assessed using population vector (PV) correlation. Union cells (neurons classified as place cells in at least one of the two compared sessions) were used to construct population activity vectors for each spatial bin. Correlation coefficients were computed between PVs across conditions over all common spatial bins. Statistical comparisons within paired conditions were performed using one-tailed Wilcoxon signed-rank tests, whereas comparisons between independent groups employed Wilcoxon rank-sum tests.

<u>Sample preparation for electron microscopy imaging</u>

Samples for SBEM were prepared as previously described (*69*). Briefly, mice were anesthetized by i.p. injection of MMB and transcardially perfused with Ringer's solution (pH 7.4), followed by 2% PFA and 2.5% glutaraldehyde (GA) in 0.1M phosphate buffer (PB, pH 7.4). The brains were further post-fixed for overnight at 4 °C in 2% PFA/2.5%GA in 0.1M PB and then washed several times with 10% sucrose in PB (0.1M). Sections (200 µm thickness) were prepared under PB (0.1 M, pH 7.4) using a vibratome (Leica VT 1000S). Sections containing the dorsal part of the hippocampus were collected and microdissected into small pieces (~1 $mm^2$). These hippocampus-containing sections were then incubated in a freshly prepared solution containing 1.5% potassium ferrocyanide in 0.1 M PB, mixed with the same volume of 4% aqueous osmium tetroxide ($OsO_4$) for 1 h at room temperature. After the incubation, the sections were washed in Milli-Q water and incubated in the thiocarbohydrazide solution (1%, filtered) at room temperature for 25 minutes. Then, the sections were placed in a 2% aqueous $OsO_4$ solution for 30 minutes at room temperature, washed, and incubated overnight in 2% aqueous uranyl acetate at 4 °C. The following day, sections were processed for dehydration in an ascending series of ethanol (70, 80, 90, and 100%), followed by acetone on ice, then gradually equilibrated with Epoxy resin (Epon 812, TAAB). Finally, sections were flatly embedded and placed in a 60°C oven for 48 hours for resin curing and polymerization.

SBEM image acquisition and post-processing

SBEM images were acquired using Teneo VS (Thermo Fisher Scientific, USA). Resin blocks prepared from each group of animals were mounted on an aluminum stub (Ted Pella, Inc., USA), glued with electrically conductive silver epoxy (Electron Microscopy Sciences). The block was trimmed, and the resin-tissue interface was exposed using a glass knife. Tissue quality was briefly confirmed by transmission electron microscope JEOL1400 Flash (JEOL, Tokyo, Japan). The block surface was coated with gold sputtering to enhance electron conductivity (VE-3030CVD, Vacuum Device, Inc., Japan) before installation in the microscope chamber. Aluminum stubs containing the specimen were placed on the SBEM stage, and images were acquired at an acceleration potential of 2.0 kV, a dwell time of 500 ns, and a z-step size of 50 nm. All specimens were imaged at 6,890× magnification, working distance was regulated within 6.5~7mm with the final pixel size being 4.5nm; covering a horizontal field width of 18.432 µm at a resolution of 4096 x 4096 pixels. Stacks of 158-582 images in the stratum radiatum of the dorsal CA1 were acquired from each mouse (2,684~9,903 $\mu m^3$). Serial images over the z-axis were automatically aligned using ImageJ/Fiji and the TrackEM2 plugin (https://imagej.net/plugins/trakem2/tutorials).

Segmentation of EM images

Two independent approaches were used for EM image segmentation. The first approach was manual segmentation. For this approach, dendrites, spines, and PSDs in the aligned images were segmented by two trained annotators (YJL and ATN) using Reconstruct software (https://synapseweb.clm.utexas.edu/software). A synapse was defined by the presence of presynaptic bouton with synaptic vesicles adjacent to a PSD at the cellular membrane facing spine. PSD was traced at the interface on the postsynaptic membrane and surface area was measured in 3D reconstructed data. Manually segmented objects were then exported as a 3D vector format (.dxf) for further quantification. Distances between synapses were computed as distances between the two closest PSD centroids for each PSD identified. The volumes of spines were computed as the 3D volumes of spine heads and necks.

The second approach was automated segmentation using an AI-assisted pipeline, CDeep3M (*40*). Manually segmented EM images from Pre-QIH and QIH groups were used to train models. Briefly, small fractions of aligned images (1024 x 1024 pixels, 20 z-stacks) were randomly selected and manually segmented to identify PSDs and pre-synaptic vesicles. The EM images and corresponding segmentation labels were first augmented for model training. Using these augmented image sets, models were repeatedly trained for 50,000 or 105,000 iterations, respectively, for PSD or vesicle segmentation. These models were used to segment PSD and vesicles for the datasets. The segmented images were further processed for quantification. The vesicle images were first Gaussian-filtered with a 10-sigma kernel to merge vesicle clusters. Then, PSD and vesicle segmentation images were filtered based on pixel intensities (100 for PSDs, 50 for vesicles) and size threshold (500 pixel$^2$ for PSDs, 200 pixel$^2$ for vesicles). The products of these processes were compared with raw EM images, and their matches were confirmed by the experimenter (KZT). Pairs of a PSD and a vesicle cluster with their centroids located closer than 70 pixel length were considered as synapses.

<u>Contextual fear conditioning</u>

The NIR Video Fear Conditioning (VFC) system (Med Associates, USA) was used for contextual fear conditioning. All mice were single-housed prior to the surgery. Mice in No QIH control group were injected with AAV-DIO-mCherry, instead of AAV-DIO-hM3Dq-mCherry, but followed the same experimental procedure, including CNO injection. During conditioning, mice were allowed to explore context A (a fear conditioning chamber inside of the sound-proof box, grid floor, ceiling light, ethanol odor, white noise) for five minutes prior to the onset of five footshocks (0.75 mA, 2 seconds, ITI = 30 seconds), then returned to their homecages 30 seconds after the last shock. The next day, contextual fear memory prior to QIH (Pre-Test) was assessed by placing the mice in contexts A and B (the same chamber, plastic flat floor, dim light, acetic acid odor, no noise, A-type ceiling) for 5 minutes. One day after Pre-Test, QIH induction was conducted as described above. After QIH session, mice were returned to their home cages. Five days after QIH, contextual fear memory was assessed by placing the mice in contexts A and B for 5 minutes (Post-Test). After Post-Test, mice were perfused to confirm hM3Dq-mCherry or mCherry expression in AVPe.

<u>Spatial memory task</u>

The Multi-Maze system (Ugo Basile, Italy) was used for a spatial memory task. All mice were single-housed prior to the surgery. Mice in No QIH control group were injected with AAV-DIO-mCherry, instead of AAV-DIO-hM3Dq-mCherry, but followed the same experimental procedure, including CNO injection. At least 1 week before the training, food deprivation started. More specifically, the animals were given 2.0 g of food pellets for 40 g of body weight each day. During the food deprivation, animal's body weight was carefully monitored. When more than a 20% reduction in body weight was observed and the total body weight was 24 g or less, the mouse was euthanized. After 1 week of food deprivation, mice were habituated to sugar pellets (20 mg each). On the first day of habituation, several sugar pellets were placed in their home cages. On the second day, mice were placed on the plus maze with sugar pellets scattered across it and allowed to explore for 5 minutes.

After habituation, mice were allowed to explore the maze for 10 minutes with sugar pellets scattered throughout. Then, mice were trained to navigate from a randomized starting arm to a fixed goal arm. Specifically, mice were placed at the beginning of the starting arm and allowed to explore for three minutes. If mice entered the correct goal arm, they were allowed to consume the sugar pellets and then removed from the maze. If mice entered the wrong arm, they were immediately removed from the maze. If mice returned to the starting arm or failed to choose another arm within 3 minutes, they were removed and scored as failures. The arm opposite the starting arm was closed. After a trial, mice had a 20-30 second inter-trial interval. When mice completed 10 correct trials or 20 total trials, they were returned to their homecages. Mice were trained daily until their performance (correct choices/correct + wrong choices) exceeded 90% for two consecutive days.

One day after completing training, QIH was induced as described above. Five days after QIH session, the post-session and probe test were conducted. The probe test followed the same procedure as the training trials, except that sugar pellets were not placed on the goal arm. The test continues until 10 correct choices or up to 20 trials. The percentage of correct choices was used as a measure of spatial memory performance.

Hippocampal lesion

Contextual fear conditioning was used to test whether contextual fear memory depends on the hippocampus after arousal from QIH. As in the previous experiment, mice injected with the AAV vector were fear-conditioned in context A and tested in contexts A and B, followed by QIH induction. One day after arousal, their hippocampus (both dorsal (AP -1.5 mm; ML ±2.0 mm; DV -2.0 mm) and ventral (AP -3.0 mm; ML ±3.0 mm; DV -3.0 mm)) received stereotaxic infusion of NMDA (N-methyl-D-aspartate) solution (0.15 μL per cite at a rate of 0.1 μL/min, 10 mg/mL PBS) to achieve an excitatory lesion. Five days after the lesion, mice were exposed to contexts A and B, and their freezing behavior was scored using Video Freeze Software (Med Associates, USA).

One-photon $Ca^{2+}$ imaging

We recorded $Ca^{2+}$ dynamics of the CA1 pyramidal cells in the dorsal hippocampus using an integrated miniature fluorescence microscope (nVista 2.0, Inscopix). Qrfp-iCre::Thy1-GCaMP6f mice were injected with either AAV-DIO-hM3Dq-mCherry and received implantation of the GRIN lens targeting the CA1 as previously described. Three weeks after surgery, their $Ca^{2+}$ activities were recorded while exploring four different open fields (context A, B, C, and D; A, white tray (32.5 cm x 44.5 cm), plastic floor, dim room light, EtOH odor; B, the white tray (32.5 cm x 44.5 cm) with four colored walls surrounding, floor covered with kim wipes, bright room light, acetic acid odor, background white noise; C, yellow rectangular box (30 cm x 40 cm) with visual cues attached on walls inside, bedding materials on floor, dim room light, benzaldehyde odor; D, deep white box (32.5 cm x 51.5 cm), floor covered with bubble wrap, bright room light, isoamyl alcohol odor). On Day 1, mice explored context A and then context B for 15 minutes each, with a 5-minute interval. During the exploration, their behaviors were recorded by a camera (acA1300-75gc GigE camera, Basler, USA) located 78.5 cm above the arena using Cheetah software (Neuralynx, USA), while $Ca^{2+}$ activity was acquired using nVista Data Acquisition Software (version 2.3.0, Inscopix, USA). Time stamps from the two acquisition systems were aligned by sending TTL pulses from Inscopix to Cheetah. Behavioral tracking was obtained by a custom MATLAB script implementing background subtraction, morphological filtering, and image inversion to generate a high-contrast mask of the animal's body, from which the centroid was extracted. A post hoc constant-velocity Kalman filter was applied to smooth the trajectory and correct faulty detections. On Day 2, the same context A and B recordings were conducted. On Day 3, QIH was induced as previously described, but without recording $Ca^{2+}$ activities. Mice were kept in the hibernation refrigerator (20 °C) for 48 hours, then returned to their home cages. On Day 10 and 11, the same context A and B recordings were conducted. After this QIH recording session was completed, the same animals were recorded for No QIH recording session. The identical experimental procedure was conducted, except for context C and D, and saline injection (instead of CNO) on Day 3. After QIH and No QIH sessions, mice were perfused, and their expression and GRIN lens locations targeting CA1 were confirmed.

Two-photon spine imaging

For all in vivo two-photon spine imaging, fluorescent images were acquired using a two-photon laser-scanning microscope (A1R-MP+, Nikon) equipped with non-descanned detectors and a GaAsP PMT. High-resolution spine images (94.3 × 94.3 $\mu m^2$; 92.1 nm/pixel) were obtained using a ×25/1.0 NA water-immersion objective lens (XLPLN25XSVMP2, Evident). The correction collar of the objective lens was adjusted by maximizing the fluorescence brightness of the acquired images. The excitation laser source was a Ti:Sapphire laser (MaiTai eHP DeepSee, Spectra Physics), tuned to a wavelength of 960 nm and running at a repetition rate of 80 MHz. The emitted fluorescence signal was separated using a 560-nm dichroic mirror. Image stacks were acquired with a Z-step interval of 1 μm. The detector sensitivity and laser power were optimized for each individual experiment to ensure an adequate signal-to-noise ratio. The laser power was typically around 5–25 mW.

Longitudinal two-photon microscopy was performed on head-fixed mice to track dendritic spine dynamics. To compare changes in spine morphology before and after QIH induction, Z-stack images of the identical dendritic regions were acquired at four distinct time points: 0d (before CNO administration), 1d, 3d, and 7d after QIH induction. To compensate for motion artifacts, all acquired image stacks were motion-corrected using the StackReg plugin in ImageJ. The corrected images were then processed using an Unsharp Mask filter (radius = 8.0, mask = 0.6) to enhance the morphological details of the dendritic spines. Following these processing steps, the spine number and spine head area were manually quantified using ImageJ.

Long anesthesia (ANE)

As a behavioral state characterized by prolonged inactivity and disrupted synaptic structures, we combined long-term anesthesia with intra-hippocampal infusion of Cytochalasin D (Tocris Bioscience, USA). More specifically, one day after the Pre-Test of the contextual fear conditioning experiment, we placed the mice in the induction chamber with air and 1.5% isoflurane provided. Once the animals were anesthetized, we lowered the isoflurane concentration to 1.25% and maintained them for 3 hours, followed by another 3 hours of isoflurane exposure at 1.0%. Throughout the anesthesia, a heat pad was provided beneath the induction chamber. After 6 hours of anesthesia, the animals were returned to their home cages. The next day, the same anesthesia treatment was performed. At the end of the second anesthesia, the mice were placed in the stereotaxic apparatus and received the intra-hippocampal infusion of Cytochalasin D (0.5 μL, 0.5 mg/mL in 2% DMSO, PBS) targeting the dorsal CA1 (AP, -2.0 mm; ML, ±1.5 mm; DV, -1.5 mm) bilaterally. Five days after the two days of anesthesia treatment, mice underwent the Post-Test by placing them in context A and B.

For tetrode recording during anesthesia, mice were placed in a familiar plastic bucket, and their spike and LFP activities were recorded for 15 minutes, as previously described. After this baseline recording without anesthesia, mice were disconnected from the recording rig and exposed to 1.5% isoflurane in the induction chamber. Once the animals were anesthetized, the Microdrive was reconnected to the Neuralynx rig and recorded for 1 hour while 1.0% of isoflurane was supplied.

For EM imaging, eGRASP, and quantification of synaptic contacts, the one day of anesthesia were conducted, followed by the intra-hippocampal infusion of Cytochalasin D. Soon after the infusion, mice were perfused for EM imaging or histology.

eGRASP labeling and engram synapse imaging

Qrfp-iCre::c-Fos-tTA mice were provided with the doxycycline-containing pellets (ON DOX). The animals were injected with either AAV-DIO-hM3Dq-mCherry or mCherry alone to AVPe, AAV-TRE-mScharlet-P2A-post-eGRASP to CA1, and AAV-TRE-yellow-pre-eGRASP to CA3. Two to three weeks after surgery, mice were switched to normal food pellets for 48 hours without disturbance (OFF DOX), followed by the contextual fear conditioning as previously described. Soon after conditioning, mice were given high-concentration doxycycline-containing pellets (1 g/kg, Bioserv) to suppress further tagging of c-Fos-positive cells immediately. The next day, the mice were either entered sham QIH in the refrigerator and then were perfused 24 hours after CNO injection (NoQIH group), entered QIH and then were perfused 24 hours after CNO injection (QIH group), or anesthetized with isoflurane for six hours and then infused in the hippocampal CA1 with Cytochalasin D (ANE group) prior to perfusion. The brains were post-fixed in 4% PFA for 24 hours and sectioned (50 µm thickness) using a vibratome (VT1000S, Leica).

Brain sections mounted with a mounting medium (ProLong Gold Antifade Mountant with DAPI, Thermo Fisher Scientific, USA) were imaged using a confocal microscope (LSM900, Zeiss). Confocal images were obtained as stacks of 40-50 planes (0.5 or 1.0 µm steps, 63x objective lens (oil-immersion, N.A./1.4), 1726 x 1726 pixel scanning resolution (pixel size = 0.06 µm)) together with DAPI images. eGRASP puncta were manually counted. eGRASP signals outside of mScarlet-labeled dendrites or inside DAPI nuclei were excluded as non-specific signals. eGRASP puncta located within 5 µm distance were counted as clustered eGRASP puncta, and the other eGRASP puncta were counted as isolated eGRASP puncta.

Correlative light and electron microscopy (CLEM)

eGRASP labeling was conducted as described above. A day after labeling and subsequent ON DOX, the animals entered QIH and then were perfused 24 hours after CNO injection. The brains were post-fixed with 2% PFA/2.5% GA in 0.1M PB for 24 hours, and sectioned (200 µm thickness) by using a vibratome (VT1000S, Leica).

Brain sections containing the dorsal hippocampus were then imaged under a confocal microscope equipped with a multiphoton laser (LSM980 Airyscan2, Zeiss) using a 40× water-immersion lens (NA, 1.2). More specifically, clustered eGRASP puncta were first identified as described above. Then, an ROI containing the clusters was marked with a smaller L-shape branding mark (10 µm length x 5 µm width) at the same z-depth using a femtosecond-pulsed titanium-sapphire laser with the combination of 800 nm with 100% laser power and a laser output of ~3,000 mW (Camereon, COHERENT), and scan duration 8.19-32.77 µs/pixel were applied. At the z-depth 2.5 µm above the target depth, another flipped L-shape branding mark was created (100~120 length x 5μm width). This second branding mark served as a landmark to initiate SBEM image acquisition. After branding marks were created, two sets of z-stack confocal images were obtained (40x water-immersion lens (NA, 1.2): 0.5 μm z-steps to cover the area between 2 branding marks with additional digital zooms (2.0x); 40x water-immersion (NA, 1.2): 1.0 μm z-steps to image the entire z-volume) to correlate the confocal and EM z-stack images.

After the confocal imaging, the brain sections were post-processed for SBEM as described above. Once the second branding mark was confirmed under the ultramicrotome (UC7, Leica), the block was coated with gold by sputtering (Ion Coater IB-3, Eiko, Japan) and installed in SBEM (Merlin, Zeiss) equipped with a 3View in-chamber ultramicrotome system, Digital Micrograph, and OnPoint detector (Gatan, Inc.). Images were acquired at an acceleration potential of 1.3~1.7kV, at 4.5 nm/pixel resolution with a dwell time of 1.0 μs/pixel and a slice thickness of 40–50 nm. 82.944 μm x 82.944 μm x 5-10 μm of z-stack images were acquired to cover sufficient ROIs containing the target clustered eGRASP signals and surrounding structures, including axonal boutons. Sequential images were processed at reduced resolution (50%) in FIJI (https://fiji.sc/) and aligned using the TrakEM2 plugin with least-squares translation and elastic alignment.

The confocal and SBEM images were correlated and analyzed manually. Briefly, multiple landmarks, such as blood vessels, cell nuclei, engram dendrites, and shapes of the branding marks, were identified, and some were manually segmented using Vast Lite (https://lichtman.rc.fas.harvard.edu/vast/). Manual segmentations were repeated until matching of multiple landmarks and engram dendrites between the confocal and SBEM images was obtained. On the matched engram dendrites, clustered eGRASP-positive spines were identified in SBEM images, and their corresponding synapses were determined. From the eGRASP positive spines, their pre-synaptic axonal boutons were further segmented manually and determined if the same bouton makes another synaptic contact (multi-synaptic bouton, MSB) or not (non-MSB). For comparison, SBEM datasets from No QIH animals were used to estimate the overall density of MSBs under an awake state. From manually segmented dendrites, spines were randomly selected, and their pre-synaptic axonal boutons were traced to determine if they made synaptic contact with another spine. The proportion of MSBs in this analysis served as the likelihood of observing MSBs if spine elimination occurred randomly during QIH. The spine volumes of the clustered eGRASP positive synapses were quantified from manual trace. The spine volumes of randomly selected spines on the same mScarlet positive dendrites were compared to determine if these clustered engram synapses have larger spines or not.

Statistical tests

Statistical analyses were performed using MATLAB (2024b, MathWorks), R, and GraphPad Prism (Version 10, GraphPad Software). Data in the main text and figures are represented as mean ± the standard error of the mean, or as boxplots indicating the median, $25^{th}$, $75^{th}$, the maximum/minimum value within 1.5x the interquartile range, and the outliers. Sample number (n) indicates the number of slices, 3D blocks, or mice in each experiment. The significance p-value was set at 0.05 or below. In each figure, p-values less than 0.05, 0.01, 0.001, 0.0001 were presented as *, **, ***, ****, respectively. The data points were first tested for normality using the Kolmogorov-Smirnov test, except for pyramidal cell firing rates, which are known to follow a log-normal distribution (*70*). For data points following the normal distribution, either a two-tailed Student t-test or ANOVA, followed by post-hoc comparisons with p-value corrections as described in the legends, was used. For data that violated normality assumptions, non-parametric tests (Mann-Whitney U test, Wilcoxon rank-sum test, or signed rank test) were used for comparisons. When appropriate, pair-wise post-hoc non-parametric rank sum comparisons were conducted with their alpha values corrected by the number of comparisons (Dunn's method). For comparisons of the proportion of MSBs in CLEM data, Fisher's exact test was used.

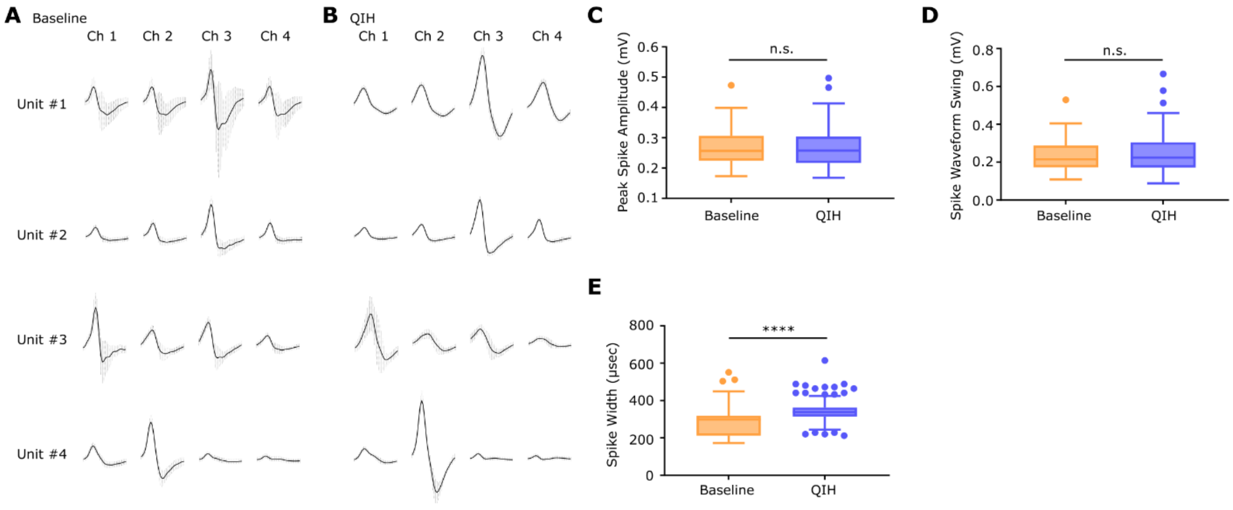


**Fig. S1. Spike waveforms are largely unchanged, except for longer spike width, during QIH.** (**A**) Representative spike waveforms (Unit #1-4) recorded from a single tetrode (Ch 1-4) in the CA1 pyramidal cell layer during the baseline recording session. Solid lines indicate the mean waveform, and dashed lines indicate the standard deviation across spikes of the same unit. (**B**) Representative spike waveforms of the same units (Unit #1-4), 4 hours after CNO injection. Same as (A), solid lines indicate the mean waveform and dashed lines indicate the standard deviation across spikes of the same unit. (**C**) Boxplots comparing peak spike amplitudes during the baseline and QIH recording sessions (n = 85 units, Wilcoxon test, p = 0.993). (**D**) Boxplots comparing spike waveform swing amplitudes during the baseline and QIH recording sessions (n = 85 units, Wilcoxon test, p = 0.4046). (**E**) Boxplots comparing spike width during the baseline and QIH recording sessions (n = 85 units, Wilcoxon test, p < 0.0001). Box plots show median, 1st and 3rd quantiles, minimum & maximum values within 1.5 times the interquantile range (IQR) from each quantile, and the other data points outside of the range.

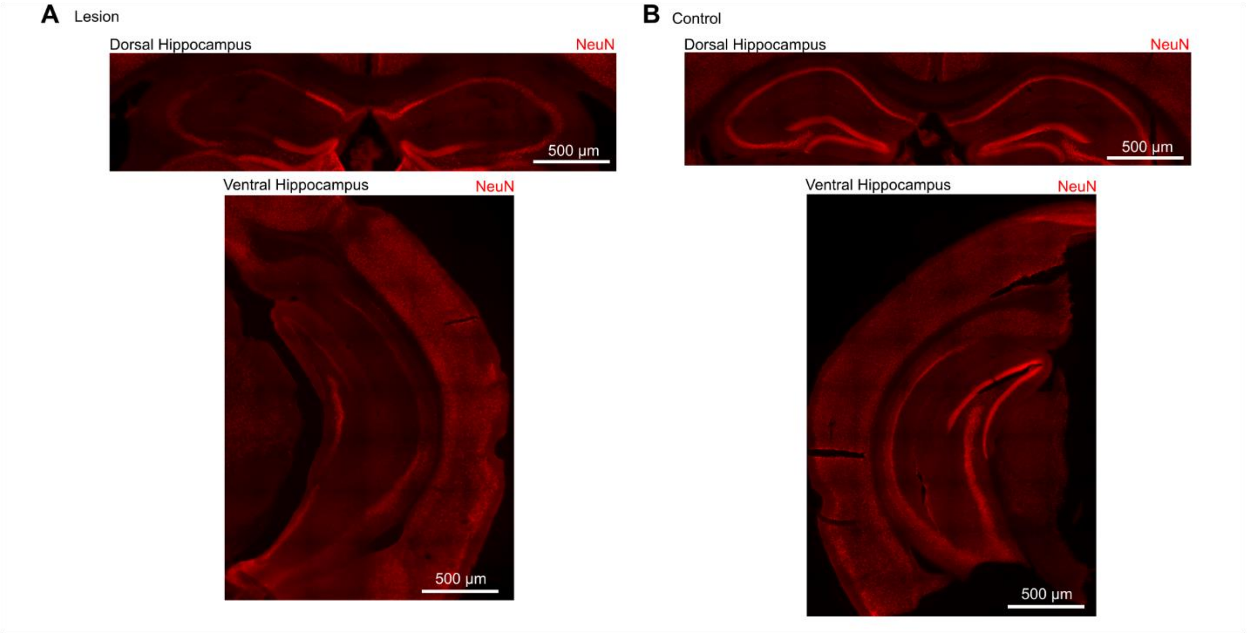


**Fig. S2. Hippocampal lesion by infusion of NMDA.** (**A**) Representative images of the coronal sections indicating the hippocampal lesion after NMDA infusion. NeuN signals are absent in the pyramidal cell layers in both the dorsal and ventral hippocampus. (**B**) Representative images of the coronal sections from a control mouse.

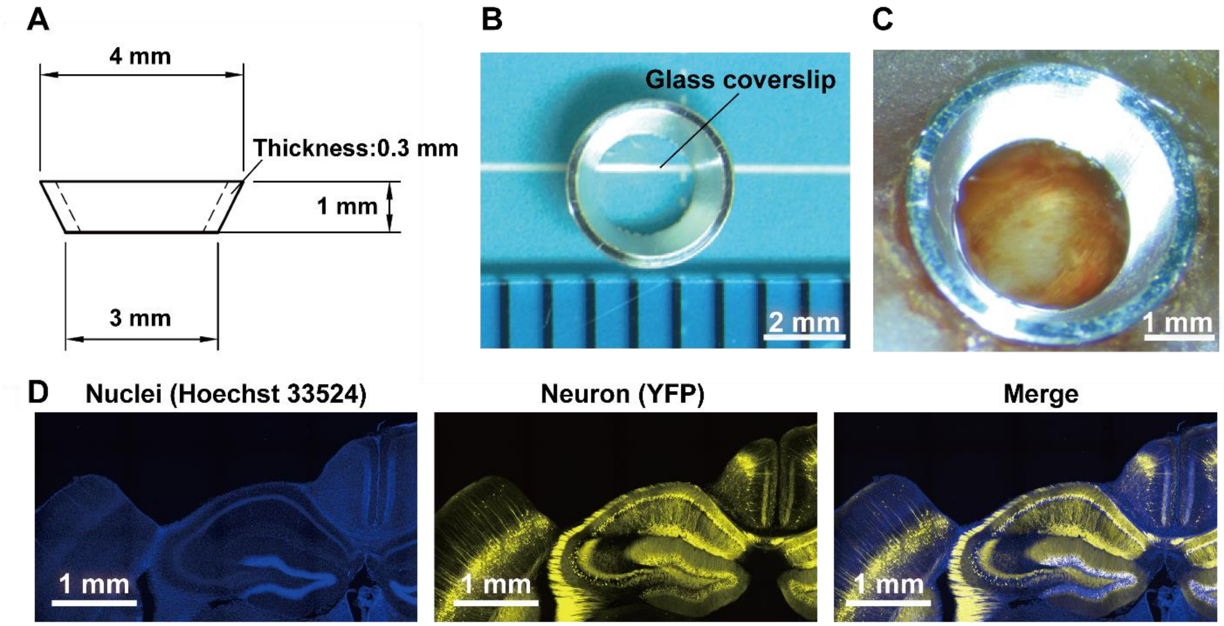


**Fig. S3. Two-photon spine imaging from the hippocampal CA1 through a custom-made cannula.** (**A**) The design of the custom-made cannula. (**B**) The image of the cannula. (**C**) The cannula was implanted in the mouse brain. Note that the hippocampal corpus callosum is exposed inside the cannula. (**D**) Schematic images indicating the surgical procedure. (E) Representative images of the coronal sections indicating YFP expression in neurons, the ROI, and cortical removal.

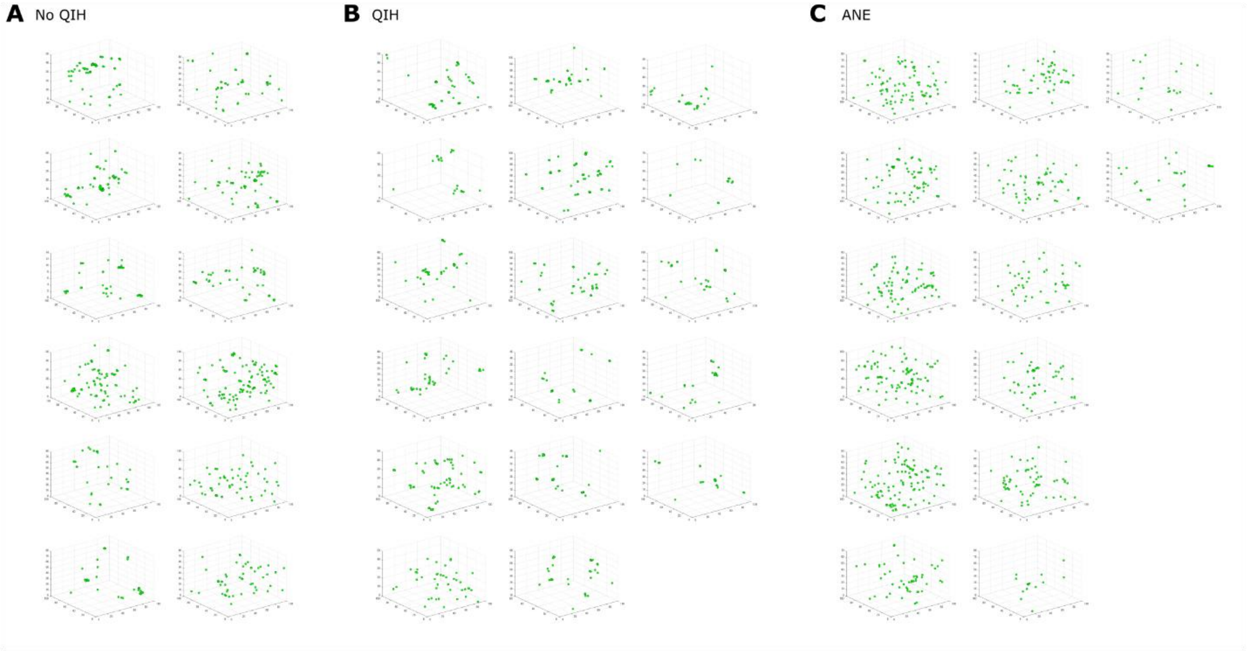


**Fig. S4. Three-dimensional projections of eGRASP signals.** (**A-C**) eGRASP puncta are plotted in a 3D space for each sample used to quantify engram-engram synapses (related to Fig. 5I). Units are micrometers. Multiple eGRASP puncta within 5 $\mu$ m vicinity are counted as a single clustered eGRASP.

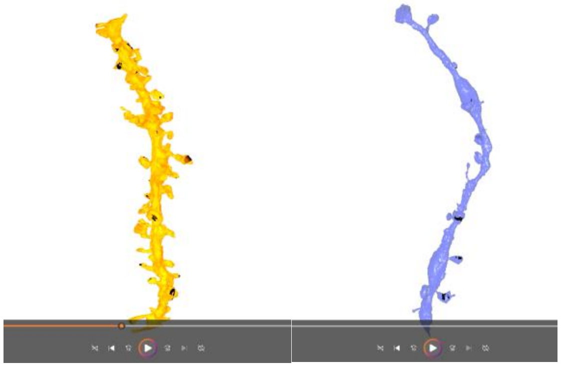

**Movie. S1. Dendritic structures reconstructed from SBEM images.** (**A**) A representative dendrite from manual segmentation of SBEM images (No QIH animal). (**B**) A representative dendrite from manual segmentation of SBEM images (QIH animal). The black shadowed areas indicate post-synaptic densities (PSDs).

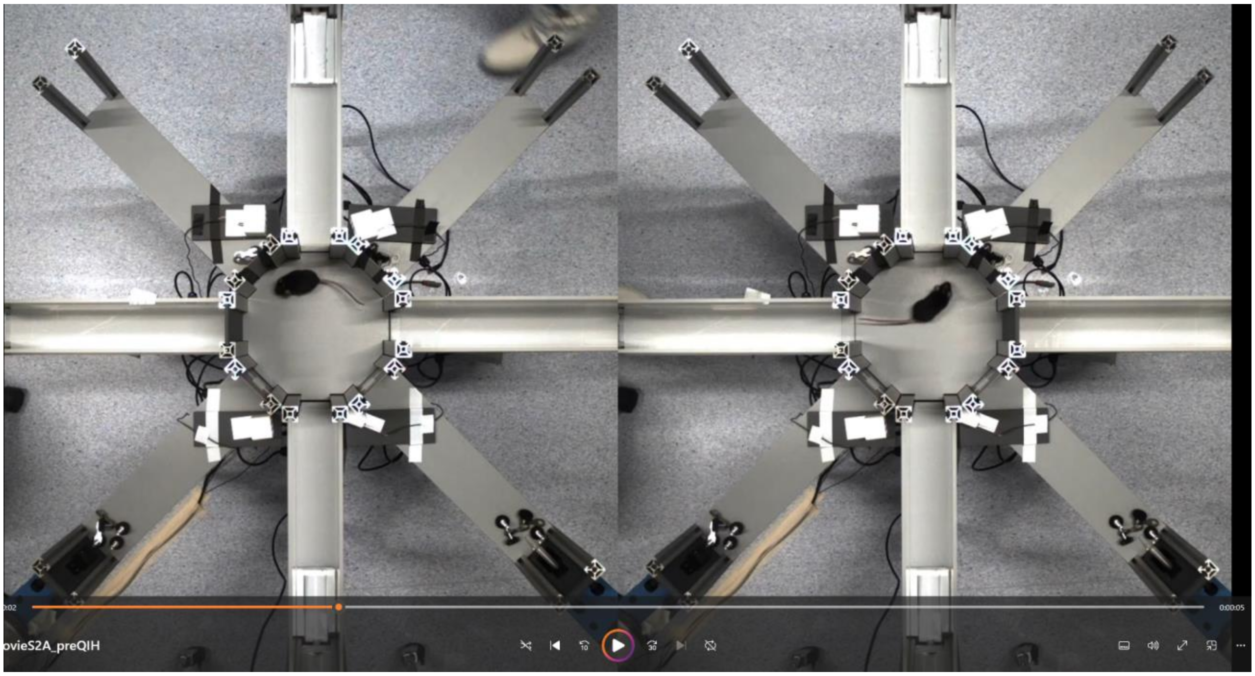


**Movie. S2. Intact spatial memory after QIH.** (**A**) Representative trials of the spatial memory task, Pre-QIH. Left, a representative trial starting from the west arm. Right, a representative trial starting from the east arm. (**B**) Representative trials of the spatial memory task, Post-QIH. Left, a representative trial starting from the west arm. Right, a representative trial starting from the east arm.
